# Motor Learning Drives Dynamic Patterns of Intermittent Myelination on Learning-activated Axons

**DOI:** 10.1101/2021.10.13.464319

**Authors:** Clara M. Bacmeister, Rongchen Huang, Lindsay A. Osso, Michael A. Thornton, Lauren Conant, Anthony Chavez, Alon Poleg-Polsky, Ethan G. Hughes

## Abstract

Myelin plasticity occurs when newly-formed and pre-existing oligodendrocytes remodel existing patterns of myelination. Recent studies show these processes occur in response to changes in neuronal activity and are required for learning and memory. However, the link between behaviorally-relevant neuronal activity and circuit-specific changes in myelination remains unknown. Using longitudinal, *in vivo* two-photon imaging and targeted labeling of learning-activated neurons, we explore how the pattern of intermittent myelination is altered on individual cortical axons during learning of a dexterous reach task. We show that behavior-induced plasticity is targeted to learning-activated axons and occurs in a staged response across cortical layers in primary motor cortex. During learning, myelin sheaths retract, lengthening nodes of Ranvier. Following learning, addition of new sheaths increases the number of continuous stretches of myelination. Computational modeling suggests these changes initially slow and subsequently increase conduction speed. Finally, we show that both the magnitude and timing of nodal and myelin dynamics correlate with behavioral improvement during learning. Thus, learning-activated, circuit-specific changes to myelination may fundamentally alter how information is transferred in neural circuits during learning.

---

In the CNS, oligodendrocytes extend numerous processes to enwrap nearby axons with discrete myelin sheaths. Myelin is traditionally known for its role in saltatory conduction, where its insulating properties accelerate action potential propagation by allowing local electrical circuits only to be generated at nodes of Ranvier^1^. However, the discovery of intermittently myelinated axons^2^, with large gaps between individual myelin sheaths, raises the possibility that sheath location and diverse patterns of myelination could represent an additional avenue through which experience-dependent plasticity could modulate signal transmission between neurons.

While myelination begins during prenatal development, this process continues throughout life and is correlated with the functional maturation of specific brain circuits^3^. Recent work shows that neuronal activity and naturalistic experiences during adulthood regulate the formation of new myelin by modulating the generation of adult-born oligodendrocytes^4,5^. Importantly, this process is required for learning novel behaviors and the consolidation of new memories^6–9^. However, the link between learning-relevant neuronal activity and circuit-specific changes in myelination remains unknown.

Recent studies show that neuronal activity can regulate myelin sheath thickness, length, and placement during oligodendrocyte generation and maturation. Exogenous stimulation (electrical, optogenetic, pharmacological, and chemogenetic) or interruption of neuronal activity biases the selection of axons targeted for myelination^10–12^. Furthermore, neuronal activity regulates elongation by modulating calcium signaling within nascent myelin sheaths^13,14^. These types of adjustments during myelin sheath generation and placement can be used to fine tune communication between neurons by modifying timing delays^15–17^ and synchronizing signals from distant areas^18^ during the functional maturation of these circuits.

Long thought to be a stable structure once generated, studies show that extrinsic signals can remodel pre-existing myelin sheaths and nodes of Ranvier outside of the initial myelin generation process. Following oligodendrocyte differentiation, the length of myelin sheaths is largely established in 3-7 days^19,20^. However, in developing zebrafish, mature myelin sheaths continue to elongate to compensate for body growth and ablation of neighboring sheaths induces remodeling of remaining sheaths^19^. Recent studies suggest that transcranial magnetic stimulation or spatial learning can modify the length of node of Ranvier, altering conduction velocity^21^. Moreover, sensory deprivation induces experience-dependent remodeling of preexisting sheaths on distinct neuronal subclasses^22^.

However, our understanding of the consequences and implications of modifying myelination as a form of neural plasticity remains incomplete. Whether myelin is modified in specific neuronal circuits during learning or life experience is entirely unknown. Furthermore, how changes in new and pre-existing myelination translate into computational alterations required for modulating behavioral output remains poorly understood. The intermittent pattern of myelination, which persists throughout life^5^, may be an anatomical anomaly in the adult CNS, or, alternatively, it may represent a distinct and dynamic design used to fine tune the function of neuronal circuits underlying learning and behavior.

Through longitudinal, *in vivo* imaging of individual oligodendrocytes, myelin sheaths, and neuronal axons in learning-activated neuronal ensembles, we show that myelin and nodal plasticity are specifically targeted to axons involved in learning a novel forelimb reach task. Computational modeling predicts that retraction of consecutive pre-existing myelin sheaths to lengthen of pre-existing nodes during learning slows conduction, while the addition of new sheaths speeds conduction in the weeks after learning. Furthermore, we find that the extent and timing of myelin dynamics is correlated to behavioral performance on a skilled reach task. Together, these results suggest that phasic shifts in the pattern of axonal myelination within learning-relevant circuits alter neuronal computations underlying modified behavioral outputs during learning.

## Results

### Learning modulates nodes of Ranvier in a staged response

Individual cortical axons are intermittently myelinated, where large stretches of exposed axon are interspersed with regions of myelination^2^. While previous work clearly demonstrates that oligodendrogenesis is altered by learning^6,20^, how these changes affect the pattern of myelin and nodes of Ranvier on individual axons remains unclear. To determine changes to the pattern of nodes of Ranvier on individual axons over time, we used longitudinal, *in vivo* two-photon imaging in the forelimb region of motor cortex of *MOBP-EGFP* transgenic mice, which express EGFP in all mature oligodendrocytes and their associated myelin sheaths in cortex^5,20^ (**Supplementary Fig. 1**). We trained 2-3 month old *MOBP-EGFP* mice in a skilled, single pellet, contralateral forelimb reach task (**Supplementary Fig. 2**) and conducted long-term *in vivo* imaging and post-hoc immunostaining. With near-infrared branding^23^, we aligned longitudinal, *in vivo* images of myelin and nodes with post-hoc immunostained axons to resolve dynamics of pre-existing and newly-generated nodes and myelin sheaths along individual axons in response to learning (**Supplementary Fig. 1,3**). Age-matched, untrained controls were exposed to the training environment without completing the behavioral task and received the same diet restriction as learning mice for the duration of the experiment.

In the cortex, intermittently myelinated axons possess three types of sheaths that are defined by their location relative to neighboring myelin: sheaths with two directly neighboring myelin sheaths, sheaths with one directly neighboring myelin sheath, and sheaths with no direct neighbors (**Fig. 1a**). In all cases, “direct neighbors” are defined as sheaths that form nodes of Ranvier (gaps of 3 microns or less between *MOBP-EGFP+* myelin sheaths) with each other. We found that 90.90±3.73% of nodes of Ranvier visualized by *in vivo* imaging contained sodium channels in post-hoc immunostaining, a value comparable to nodes of Ranvier defined histologically in fixed tissue (Student’s t test; t(2.14) = −1.88, p = 0.19; **Supplementary Fig. 3**). Pre-existing sheaths showed three different behaviors, maintaining their length for the duration of the study (stable), lengthening over time (growing), or shortening over time (retracting). We found that the starting length of dynamic pre-existing sheaths did not differ across sheath type and initiation of sheath remodeling was distributed across the two-month imaging period suggesting that sheath age nor length factor into sheath remodeling (**Supplementary Fig. 4**). While we found growing sheaths across all timepoints, only sheath retraction was altered in learning mice relative to untrained mice (**Supplementary Fig. 4**). Next, we examined the dynamics of the three types of sheaths on intermittently myelinated axons (**Fig. 1a**). We found that learning mice have less stable sheaths across all sheath types (sheaths with 2, 1, and 0 neighbors) compared to untrained mice (**Supplementary Fig. 4**). However, when we examined specific sheath behaviors across time, we found that only retraction of pre-existing myelin sheaths was specifically regulated by learning in sheaths with either one or two neighbors (**Fig. 1b,c**). Retraction of sheaths with two neighbors was protracted and continued for two weeks after learning (**Fig. 1b**), in contrast to the retraction of sheaths with one neighbor, which occurred within the week of task acquisition (**Fig. 1c**). The growth and retraction of sheaths without neighbors was not regulated by learning (**Fig. 1d**). Therefore, learning specifically modulates the retraction of continuous stretches of myelin—that is, areas in which myelin sheaths form nodes with one or two directly neighboring sheaths. In contrast, in regions of where sheaths do not have any direct neighbors and therefore do not have any associated nodes, sheath retraction is not specifically modulated by learning.

**Figure 1.**
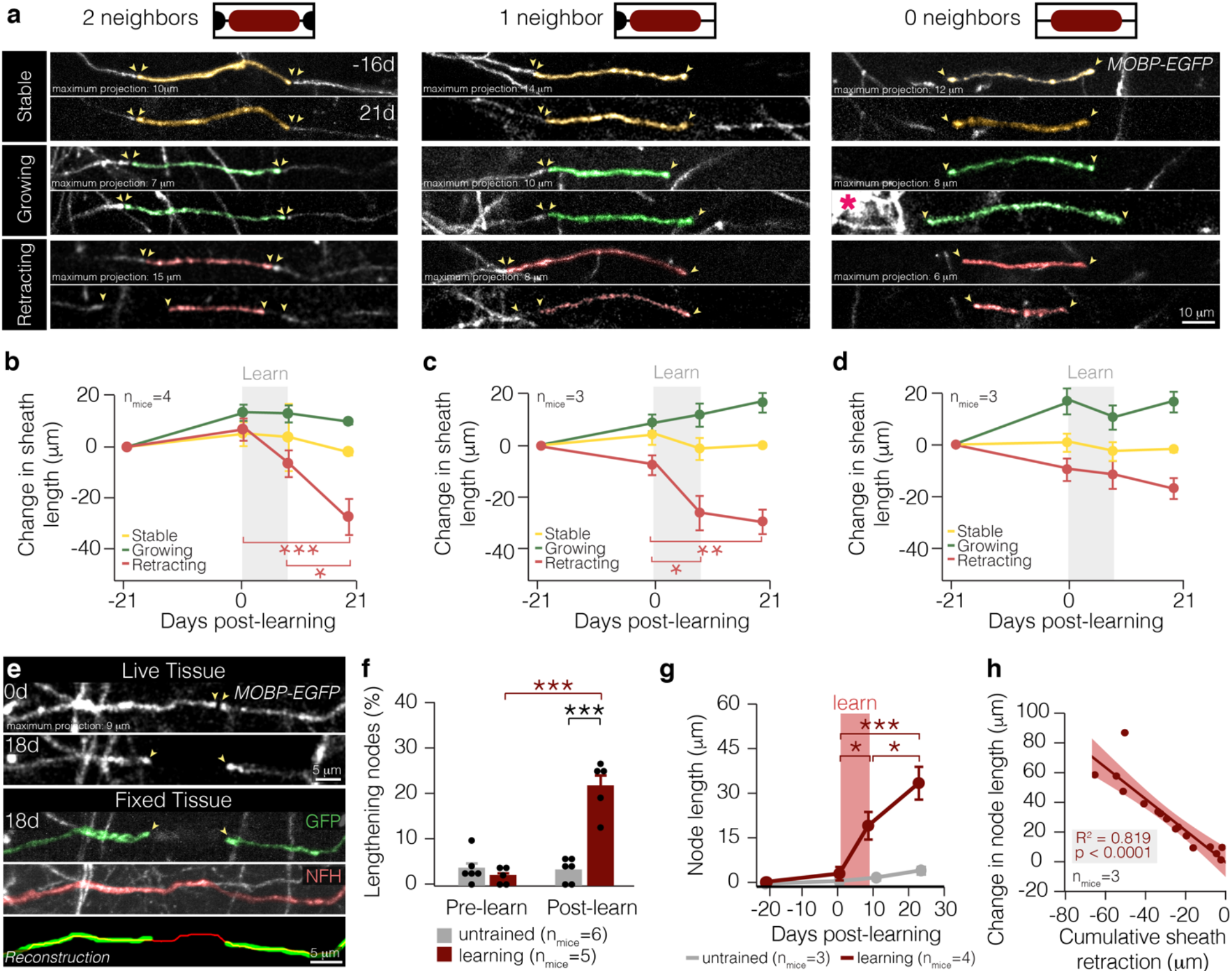
Learning-induced sheath retraction modifies node length. **a,** All types of sheaths exhibit three behaviors: stable (top), growing (middle), and retracting (bottom). Note newly-generated oligodendrocyte (middle-right pink asterisk). **b,** Retraction of sheaths with two neighbors is specifically modulated by learning (F_3,48_ = 9.04, p < 0.0001) and continues for two weeks after learning (0d vs. 21d, p < 0.0001; 9d vs. 21d, p = 0.021; Tukey’s HSD). **c,** Retraction of sheaths with one neighbor is specifically modulated by learning (F_3,40_ = 9.93, p < 0.0001) and finishes within the learning period (p = 0.029; Tukey’s HSD). **d,** Dynamics of sheaths with 0 neighbors are not specifically modulated by learning. **e**, A longitudinally-imaged, lengthened node before (0d) and after learning (18d), and that same node in fixed and sectioned tissue stained for GFP (green) and neurofilament (NFH, red). The bottom panel shows a reconstruction of the lengthened node and axon. Images manually resliced for clarity. Sheaths of interest pseudo-colored. **f,** Learning modulates proportion of nodes lengthening (F_1,9_ = 38.85, p = 0.0002). More nodes lengthen in the two weeks following learning than before learning in mice that receive motor learning (p < 0.0001; Tukey’s HSD). In the two weeks following learning, more nodes lengthen in learning mice than in untrained mice (p < 0.0001; Tukey’s HSD). **g,** Learning modulates node length (F_3,83.38_=4.88, p=0.0036). Nodes increase significantly in length during learning (p = 0.039; Tukey’s HSD) and continue to lengthen following learning (p = 0.03; Tukey’s HSD). **h,** Changes in node length are significantly correlated to changes in sheath length of associated myelin sheaths (p < 0.0001). *p < 0.05, **p < 0.01, ***p < 0.0001, NS, not significant; bars and error bars represent mean ± s.e.m. For detailed statistics, see **Supplementary Table 3, Figure 1**.

Since pre-existing, continuous myelin sheaths respond to learning by retracting, we reasoned that the length of their associated nodes may also be modulated. To explore this possibility, we conducted post-hoc immunohistochemistry on myelin sheaths, nodes, and their associated axons in mice that received longitudinal *in vivo* two-photon imaging (**Fig. 1e; Supplementary Fig. 1,3**). When we examined node dynamics in response to learning, we found that 21.73±1.61% of preexisting nodes of Ranvier elongated after learning, whereas we did not find heightened node lengthening in untrained mice (**Fig. 1f**). While both untrained and learning mice possessed nodes and gaps in myelin that shortened across the two months of imaging, learning did not modulate this parameter (**Supplementary Fig. 4)**. These learning-induced sheath dynamics resulted in an average increase in node length to 18.63±4.62 microns during reach task acquisition. In mice that engaged in motor learning, nodes continued to lengthen in the two weeks following learning, causing an additional increase in length of 14.31±5.50 microns (**Fig. 1g**). Changes to node length were highly correlated to the changes in myelin sheath length (**Fig. 1h**), indicating that sheath retraction plays an integral role in node length modification.

Learning exposed regions of previously myelinated axon via sheath retraction, however, pre-existing sheaths did not grow to add myelin to regions of exposed axon. To explore how learning might increase myelination of cortical axons, we examined how sheath addition by newly-generated oligodendrocytes changed the myelination pattern in the cortex in the weeks following learning. Learning did not affect the number of sheaths per individual newly-generated oligodendrocyte nor the length of these sheaths (**Fig. 2a-c**). However, we found that learning influenced where newly-generated oligodendrocytes placed their myelin. In untrained mice, new oligodendrocytes generated primarily sheaths without neighbors, however, after learning, new oligodendrocytes generated an increased number of continuous and fewer isolated myelin sheaths (**Fig. 2a,d**). Sheath addition after learning was more likely to be added to partially myelinated regions of an axon, filling in gaps in pre-existing myelin to create continuous stretches of myelination (**Fig. 2e,f**). These newly-filled gaps in myelination were not nodes that lengthened in response to learning. Instead, new sheaths filled in large gaps in the pattern of pre-existing myelin that were present before learning occurred (**Fig. 2g**). In 6/18 of these cases, multiple new sheaths were placed consecutively to fill in a gap in pre-existing myelination pattern (**Fig. 2h**). To further explore this phenomenon on an individual axon basis, we used longitudinal, *in vivo* imaging of *MOBP-EGFPR26-lsl-tdTomato* reporter mice injected with a virus to sparsely label local motor cortical neurons and their axons (**Supplementary Fig. 5**), which enabled us to trace axons traveling on the axis perpendicular to the imaging plane, in addition to those running parallel to the plane as done previously with our post-hoc immunostaining protocol. While lengthened nodes were not filled in by new myelin sheaths, a larger proportion of axons possess myelin and nodal dynamics in learning versus untrained mice (**Fig. 2i**). Furthermore, 27.6±7.6% of all myelinated axons in learning mice showed both node lengthening during learning and sheath addition following learning compared to zero axons in untrained mice. The majority of axons that demonstrated both node lengthening and sheath addition increased in myelin coverage following learning (**Fig. 2j**), suggesting that sheath addition was not a homeostatic response to loss of myelination due to sheath retraction and node lengthening. Overall, we found that initiation of learning-induced sheath retraction preceded the addition of new myelin sheaths in response to learning by 5.93±2.41 days (**Fig. 2k**). Together, these results indicate that motor learning-induced myelin plasticity proceeds in a phasic response in forelimb motor cortex, first by causing retraction of pre-existing myelin sheaths during learning followed by the addition of new sheaths to form continuous stretches of myelination after learning.

**Figure 2.**
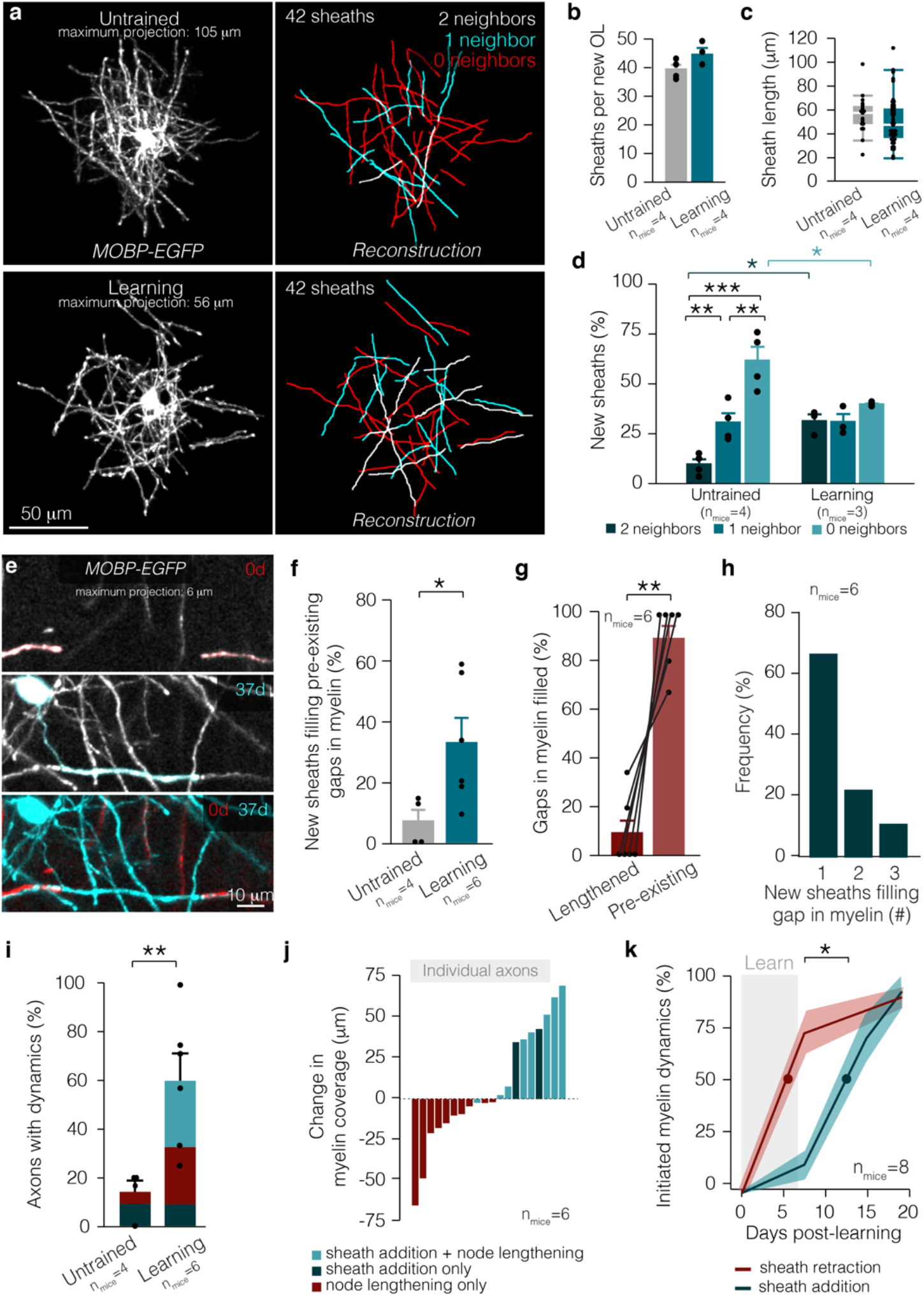
Myelin and nodal plasticity occur in distinct phases in response to learning. **a,** An entire arbor of an individual oligodendrocyte *in vivo* (left) and in reconstruction (right), in untrained (top) and learning (bottom) *MOBP-EGFP* mice. For clarity, only sheaths belonging to the relevant oligodendrocyte are shown in these images. **b,** No difference in sheath number per new oligodendrocyte in untrained and learning mice. **c,** No difference in sheath length on new oligodendrocytes in untrained and learning mice. Bars and error bars represent median and I.Q.R. **d,** Learning modulates the types of sheaths produced by new oligodendrocytes (F_5,15_=15.03, p<0.0001). Continuous sheath production is heightened after learning (p=0.048; Tukey’s HSD), while new oligodendrocytes produce fewer isolated sheaths (p=0.044; Tukey’s HSD). **e,** A new sheath (blue, 37d) fills in a gap in preexisting myelination (red, 0d) after learning (37d). **f,** After learning, more sheaths per new oligodendrocyte fill in pre-existing gaps in myelination (Student’s t-test, t(6.85)=-2.78, p=0.028). **g,** Following learning, the majority of newly filled gaps in myelination are preexisting gaps in myelination rather than nodes that lengthened (Paired Student’s t-test, t(5)=6.99, p=0.0009). **h,** The majority of pre-existing gaps in myelination are filled in by one sheath, but up to three new sheaths can be responsible for filling in a gap in pre-existing myelination. **i,** A larger proportion of axons possess myelin dynamics in learning mice (Student’s t-test, t(6.58)=-3.73, p=0.0082). **j,** Cumulative change in myelin coverage on individual axons from baseline to two weeks post-learning separated by myelin and node dynamics. **k,** Learning-induced sheath retraction precedes the postlearning burst in oligodendrogenesis and new sheath addition (Student’s t-test, t(13.43)=-2.46, p = 0.028). *p<0.05, **p<0.01, ***p<0.0001, NS, not significant; bars and error bars represent mean±s.e.m. Sheaths of interest pseudocolored. For detailed statistics, see **Supplementary Table 3, Figure 2**.

### Axonal segments exhibit synchronized nodal dynamics in response to learning

We next sought to determine whether learning-induced changes to myelin and nodal architecture were driven by signals intrinsic or extrinsic to oligodendrocytes (**Fig. 3**). We found that remodeling of myelin sheaths was not specific to individual oligodendrocytes; in fact, the majority of oligodendrocytes in both the untrained and learning conditions possessed stable, retracting, and growing sheaths (14/17 oligodendrocytes; **Fig. 3b**). In contrast, learning-induced changes in nodal length were concentrated on individual regions of axons, with changes to node length occurring at multiple nodes along the same axonal segment (**Fig. 3a,c**). In untrained mice, lengthening nodes were distributed across the population of axonal segments and the majority of nodes along individual axon regions remained stable. In contrast, individual axonal segments in learning mice had lengthening nodes that were concentrated along their length constituting the majority of their nodes (**Fig. 3c**). These findings indicate that nodes are modified in concert across select stretches of continuous myelin on individual axons in response to learning.

**Figure 3.**
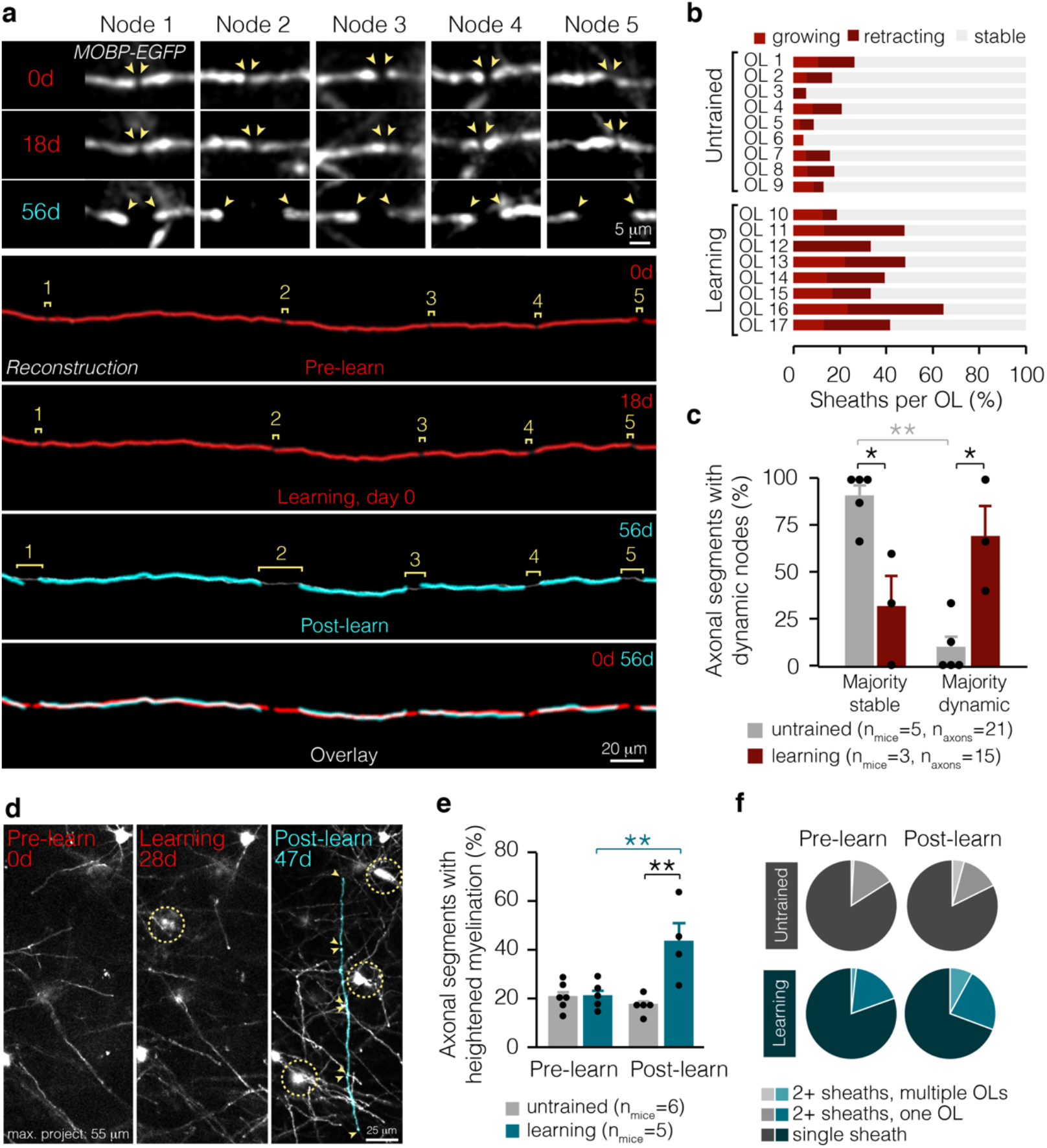
Learning-induced changes to myelin occur at specific locations along axons. **a,** Multiple nodes along the same axonal segment lengthen in response to learning *(in vivo* data shown on top, reconstructions shown on bottom). **b,** Individual oligodendrocytes in learning and untrained mice exhibit multiple sheath behaviors. **c,** Learning modulates the proportion of axonal segments with majority dynamic nodes (F_3,12_ = 14.17, p = 0.0003). In learning mice, significantly more axonal segments with dynamic myelin possess majority dynamic nodes than in untrained mice (p=0.01; Tukey’s HSD). **d,** Axons can be myelinated by multiple new sheaths. Sheaths of interest are pseudo-colored. **e,** Learning modulates the proportion of axonal segments receiving multiple new myelin sheaths (F_1,9.46_ = 14.17, p = 0.0027). In the two weeks following learning, axons are more often myelinated by multiple new sheaths than before learning (p = 0.0055; Tukey’s HSD). The percent of axonal segments receiving increased myelin sheaths is heightened in learning compared to untrained mice (p = 0.006; Tukey’s HSD). **f,** Proportion of axonal segments receiving new myelin from multiple oligodendrocytes, multiple sheaths from a single oligodendrocyte, or only one new sheath in untrained mice (gray) and before and after learning (blue). *p<0.05, **p < 0.01, ***p < 0.0001, NS, not significant; bars and error bars represent mean ± s.e.m. For detailed statistics, see **Supplementary Table 3, Figure 3.**

To determine whether sheath addition was similarly targeted to specific axonal regions after learning, we examined the placement of myelin sheaths from newly-generated oligodendrocytes. Before learning, new oligodendrocytes myelinated similar numbers of axons as in untrained mice, however, after learning, new oligodendrocytes placed more myelin sheaths on the same axonal segments in trained mice compared to untrained mice (**Fig. 3d,e**). These axon segments could receive multiple, consecutive sheaths from both individual oligodendrocytes and from more than one oligodendrocyte (**Fig. 3f**). Overall, we find that targeted sheath retraction generates new patterns of intermittent myelination during learning. After learning, we find the addition of new sheaths to pre-existing unmyelinated gaps heightens the formation of continuous stretches of myelination. Together, these results indicate that motor learning specifically modulates patterns of intermittent myelination on select axonal segments in the primary motor cortex.

### Learning induces myelin and nodal dynamics broadly across cortical layers

Previously, we showed that motor learning induced layer-specific changes in oligodendrogenesis in the primary motor cortex^20^. Our current data replicated this finding as oligodendrogenesis increases in Layer 1 (L1) but not layer 2/3 (L2/3) following learning the forelimb reach task (L1 of untrained vs. learning mice, 0.72±0.02 vs. 1.35±0.09%, p = 0.016; L2/3 of untrained vs. learning mice, 0.17±0.07 vs. 0.29±0.15%, p = 0.26). To examine whether these layer-specific differences are paired with axon-specific effects on myelin and nodal plasticity following learning, we analyzed pre-existing nodes and new sheath addition in layers 1-3 in the primary forelimb motor cortex. In L1, on mostly continuously myelinated axons, we found an increased proportion of lengthening nodes after learning, relative to node dynamics before learning and relative to nodes in untrained mice (**Fig. 4a,b**). Node lengthening occurred in concert along the length of individual axons in mice that learned the forelimb reach task compared to untrained mice (**Fig. 4c**). Similarly, the proportion of individual axons myelinated by multiple new sheaths was increased following learning in L1 (**Fig. 4a,d**). In L1, learning-induced node lengthening began during learning and preceded the addition of new myelin in response to learning by ~4 days (4.27±1.08 days) (**Fig. 4e**).

**Figure 4.**
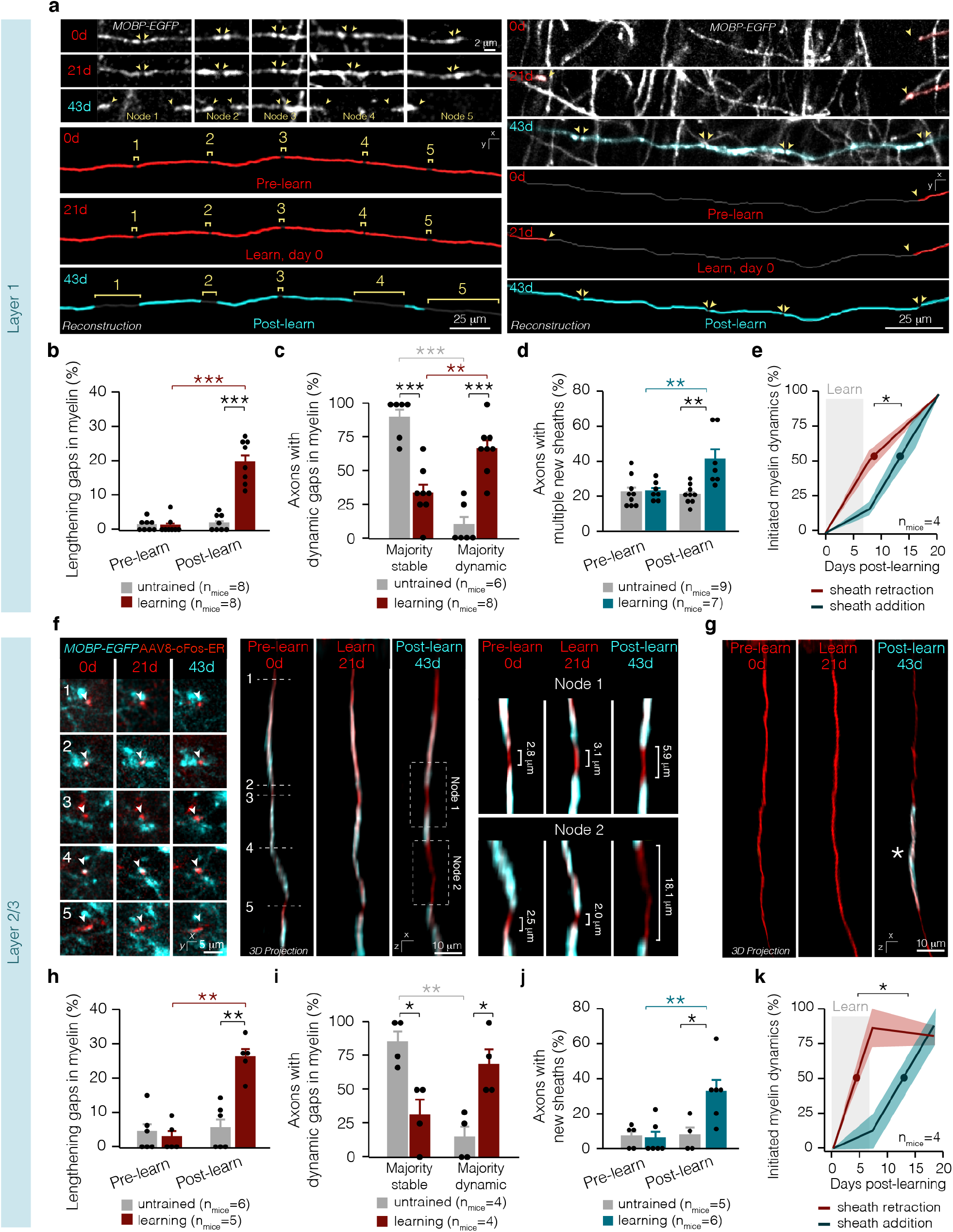
Learning induces myelin and nodal dynamics broadly across cortical layers. **a,** Max projections (top) and reconstructions (bottom) of axons with multiple nodes lengthening (left) and heightened sheath addition (right) following learning in L1. **b,** Learning modulates node lengthening in L1 (F_1,1 4_=49.33, p<0.0001). Two weeks after learning, more nodes lengthen than before learning in L1 (p<0.0001; Tukey’s HSD) and compared to untrained mice in L1 (p<0.0001; Tukey’s HSD. and L2/3 (p = 0.0014; Tukey’s HSD). **c,** Learning modulates the proportion of axons with multiple dynamic nodes in L1 (F_3,24_=25.12, p<0.0001). In learning mice, significantly more axons with dynamic myelin possess multiple dynamic nodes than in untrained mice in L1 (p<0.0001; Tukey’s HSD). In L1, learning mice have significantly more axons with multiple dynamic nodes than a single dynamic node (p=0.0052; Tukey’s HSD). **d,** Learning modulates the proportion of axons receiving multiple new myelin sheaths in L1 (F_1,11.95_=12.65, p=0.0040). Two weeks after learning, more axons receive multiple new myelin sheaths relative to before learning (p=0.0065; Tukey’s HSD) (**d**) and relative to axons in untrained mice (p=0.0087; Tukey’s HSD). **e,** Learning-induced sheath retraction precedes the post-learning sheath addition in L1 (Student’s t-test, t(4.00)=-4.51, p=0.011). **f,** Raw data in XY plane corresponding to the same axon in 3D projections (left) and reconstructions (right) with multiple nodes lengthening. **g,** 3D projection of an axon with sheath addition following learning in L2/3. **h,** Learning modulates node lengthening in L2/3 (F_1,9_=15.85, p=0.0038). More nodes lengthen after than before learning in L2/3 (p=0.0014; Tukey’s HSD) and compared to untrained mice in L2/3 (p=0.0009; Tukey’s HSD). **i,** Learning modulates the proportion of axons with multiple dynamic nodes in L2/3 (F_3,12_=9.87, p=0.0015). In learning mice, significantly more axons with dynamic myelin possess multiple dynamic nodes than in untrained mice in L2/3 (p=0.015; Tukey’s HSD) in the two weeks after learning. **j,** Learning modulates the proportion of axons receiving heightened myelination in L2/3 (F_1,8.92_=7.87, p=0.021). Two weeks after learning, more axons receive new myelin sheaths relative to before learning (p=0.0063) and relative to axons in untrained mice (p=0.048; Tukey’s HSD). **jk** Learning-induced sheath retraction precedes the post-learning sheath addition in L2/3 (Student’s t-test, t(4.8)=-2.91, p=0.035). *p<0.05, **p<0.01, ***p<0.0001, NS, not significant; bars and error bars represent mean ± s.e.m. For detailed statistics, see **Supplementary Table 3, Figure 4.**

Due to the sparse myelination of L2/3 axons^2^, nodes of Ranvier between continuous myelin sheaths were relatively rare in L2/3 compared to L1 (39.5 vs. 76.5% of sheaths forming nodes respectively). Therefore, to analyze sheath retraction, we analyzed whether the distance between myelin sheaths (rather than nodes) was altered by motor learning. Like in L1, pre-existing gaps in myelin in L2/3 lengthened in response to learning, exposing large areas of the previously myelinated axonal membrane (**Fig. 4f,g**). Similar to changes on the more heavily myelinated L1 axons, unmyelinated gaps along the length of the same axon in L2/3 lengthened in concert in learning mice, whereas axons in untrained mice had largely stable myelin profiles (**Fig. 4h**). Furthermore, while the rate of oligodendrogenesis was unaltered by learning in L2/3, we found that the proportion of axons receiving multiple new myelin sheaths was increased following learning indicating that targeted placement of newly-generated myelin sheaths is independent of the rate new oligodendrocyte formation (**Fig. 4g,j**). In L2/3, the majority of sheath retraction was initiated during learning and preceded the addition of new myelin in response to learning by ~7.5 days (7.58 ± 2.61 days (**Fig. 4j**). The delay between sheath retraction and sheath addition in L2/3 was not statistically different from the delay in L1 (Student’s t test; t(3.84) = 1.16, p = 0.311). Together, these results show that similar dynamics of targeted sheath retraction, node lengthening, and sheath addition occur across layers 1-3 of superficial cortex in response to learning in spite of varying levels of myelination and axons from distinct neuronal class subtypes. Furthermore, these findings suggest that motor learning can drive changes in myelination irrespective of changes in rates of oligodendrogenesis.

### Predicted effect of learning-induced nodal dynamics on action potential propagation

Recent evidence indicates that changes to nodal and myelin patterning can alter conduction velocity^16,17,21,24^. To explore the functional implications of axon-specific myelin and nodal plasticity during motor learning, we used the NEURON platform to simulate action potential propagation in a multicompartmental model of a myelinated axon. It is well established that the removal of myelin in pathological conditions results in the redistribution of voltage-gated ion channels into the previously myelinated axonal regions and can result in conduction failure^25–27^. Previous work shows that spike propagation in demyelinated regions depends on axonal excitability, specifically the balance of sodium:leak conductance^28^. Therefore, to test the effect of learning-induced node lengthening on axonal function, we examined the full range of potential sodium and leak conductances (gNa and gL, respectively) along newly-exposed unmyelinated regions. We examined three lengths of nodes: 1,20, 35 microns (the average length of dynamic nodes pre-learning, directly following learning, two weeks post-learning, respectively) and the effect of a large gap in myelination of 184 microns (the largest empirically observed exposed axon filled by new myelin) (see **Fig.1, Supplementary Fig. 6**, see **Methods, Supplementary Table 1** for a detailed summary of model and parameters).

In the pre-learning conditions, action potential conduction speed was greater than 2 m/s. As nodes lengthened, conduction speeds slowed, and failures became more prominent (**Fig. 5a-c**). We categorized the propagation across the unmyelinated gap in one of three ways: conduction speed above 2 m/s (“Success”), slower conduction speed (“Delayed”), and failure to propagate (“Failure”) (**Fig. 5a**). Strikingly, modification of node lengths to 20 microns resulted in propagation delays in more than 75% of the examined gNa: gL ratios (**Fig. 5c,d**). Correspondingly, the vast proportion of spikes were delayed at larger node lengths (~88% of spikes), whereas a small proportion failed to propagate (~6% of spikes) at a node width of 35 microns (**Fig. 5c,d**). These results imply that sheath retraction, node lengthening, and modulation of ion channel distributions along newly-lengthened nodes is an effective way to alter conduction velocity during learning.

**Figure 5.**
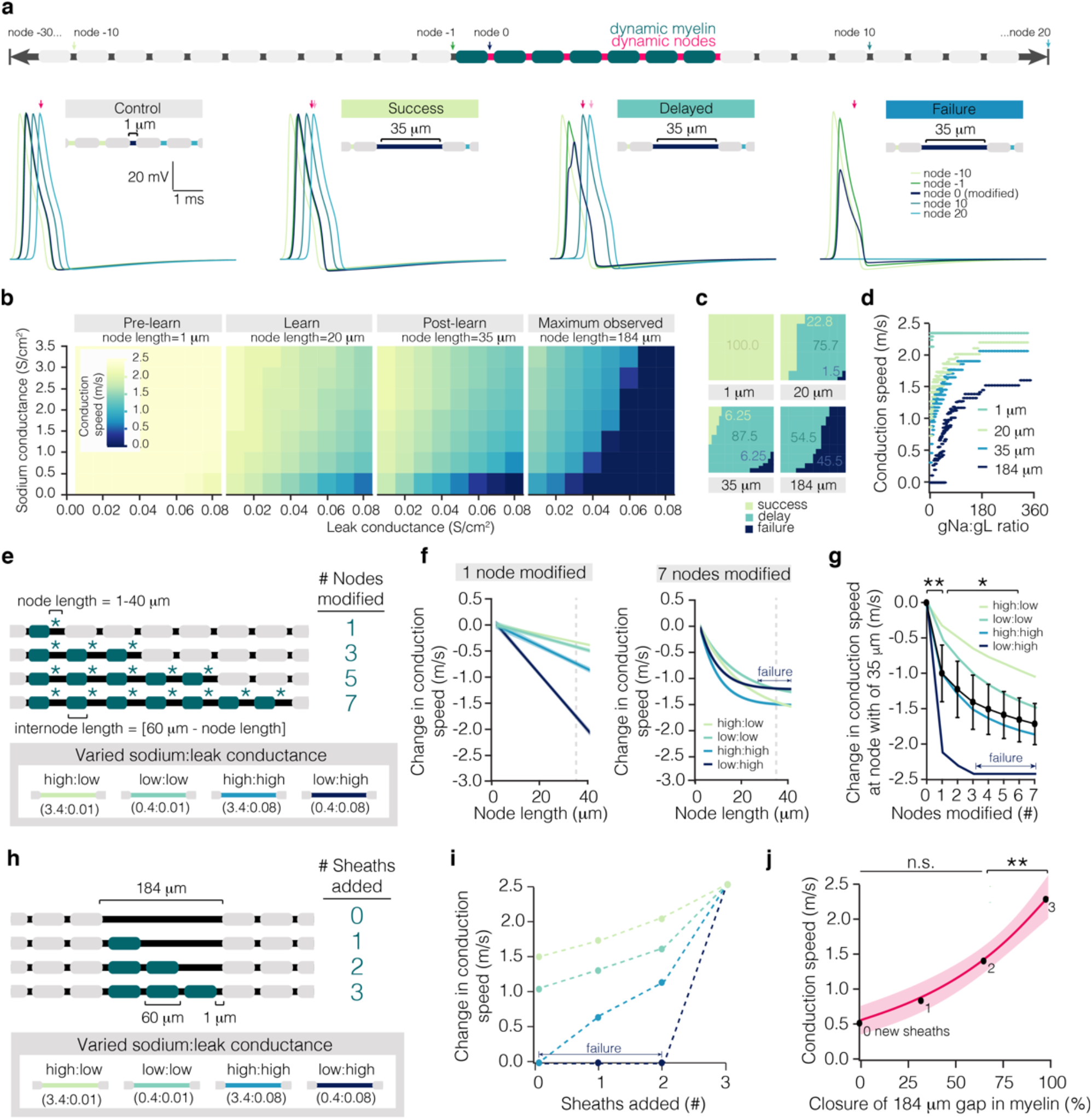
Modeled effect of learning-induced sheath and nodal dynamics on conduction. **a**, Schematic of myelinated axon model. Control trace and example traces of three categories of propagation along the modified region of axon: no change in conduction speed (“Success”), slowed conduction speed (“Delayed”), and failure to propagate (“Failure”). Note the arrival time of the control spike (dark pink) vs. the arrival time in each condition indicated (light pink). **b,** Matrix of conduction speeds across node relative to the ratio of sodium:leak conductance. Node lengths are characteristic of lengths before learning (“Pre-learn”, 1 micron), directly after learning (“Learn”, 20 microns), two weeks following learning (“Post-learn”, 35 microns), and of a large unmyelinated gap filled in by sheath addition (“Maximum observed”, 184 microns). **c,** Proportion of events that result in successful (lime), delayed (teal), or failures (navy) in propagation at 1, 20, 35, and 184 micron lengths. **d,** Modeled conduction speed as a function of sodium:leak conductance ratio at 1, 20, 35, and 184 micron lengths. **e,** Schematization of model used to predict the effect of coupled node lengthening and sheath retraction on conduction speed. Nodes were lengthened incrementally from 1 to 40 microns and associated sheaths were retracted a reciprocal amount. Asterisks indicate node retraction, while blue sheaths indicate sheath retraction. Nodes and sheaths were modified sequentially until 7 adjacent nodes were being lengthened at once. Four conductance ratios were modeled separately, using either 0.4 S/cm^2^ or 3.4 S/cm^2^ for sodium conductance (gNa) and either 0.01 S/cm^2^ or 0.08S/cm^2^ for leak conductance (gL). **f,** Modeled change in conduction speed as a function of node length at each of the four gNa:gL ratios with 1 (left) or 7 (right) nodes modified. **g,** Modeled conduction speed at a node length of 35 microns (dotted line in (f)) as a function of number of nodes modified. Learning modulates modeled conduction speed (F_7,21_ = 17.33, p < 0.0001). Modification of 1 node significantly reduces conduction speed (0 vs. 1 nodes modified, p = 0.0008; Tukey’s HSD), as does modification of 6 nodes (1 vs. 6 nodes modified, p = 0.042; Tukey’s HSD). **h**, Schematization of model used to predict the effect of adjacent sheath addition on conduction velocity. Sheaths of 60 microns in length were added sequentially to a 184 micron unmyelinated gap. **i,** Modeled change in conduction speed as a function of sheaths added. **j,** Modeled conduction speed as a function of coverage of the unmyelinated gap, averaged across the four sodium:leak conditions. Sheath addition modulates conduction speed (F_3,9_ = 14.39, p = 0.0009). The addition of a sheath that completely closes the unmyelinated gap results in a significant increase in conduction speed (p = 0.01). *p < 0.05, **p < 0.01, ***p < 0.0001, NS, not significant; bars and error bars represent mean ± s.e.m. For detailed statistics, see **Supplementary Table 3, Figure 5**.

Experimental observations revealed that up to 7 adjacent nodes could be modified in mice engaged in motor learning (see **Fig. 3c**, **Supplementary Fig. 6**). To test the effect of such numerous nodal dynamics on conduction speed, we simulated the lengthening of adjacent nodes by retracting their associated sheaths (**Fig. 5e; Supplementary Fig. 6**). We focused on 4 separate gNa: gL conductance ratios (0.4 S/cm^2^ or 3.4 S/cm^2^ for sodium conductance; 0.01 S/cm^2^ or 0.08 S/cm^2^ for leak conductance; **Fig. 5e**). When seven consecutive nodes were lengthened in concert, conduction velocity was dramatically decreased, eventually leading to failure of propagation at nodes with low excitability (**Fig. 5f**). While conduction speed decreased as more consecutive nodes were lengthened, the most significant change in propagation was introduced by modification of a single node (**Fig. 5g; Supplementary Fig. 6**). Together, these findings suggest that action potential propagation is highly tuned to the lengthening of individual nodes along the axon, with unavoidable spike delays when several nodes lengthen in response to learning.

Sheath addition following learning led to the closure of gaps in preexisting myelin sheaths, generating new stretches of continuous myelination (see **Figs. 2,3**). To determine the effect of sheath addition on conduction velocity, we modeled consecutive sheath addition into an unmyelinated gap until the gap was eliminated (**Fig. 5h**). In this scenario, we examined the largest span of an empirically observed unmyelinated gap to be completely filled in with myelin (184 microns) (**Supplementary Fig. 6**). We found that up to three sheaths can be added to a single unmyelinated gap (**Supplementary Fig. 6**), therefore, we used these parameters to model the closure of gaps in myelination. We chose a sheath value of 60 microns as this matched the average sheath length seen for continuous sheaths (**Supplementary Fig. 6**). Adding fewer, longer sheaths within the range of observed sheath lengths (**Fig. 2c**) did not significantly affect conduction speed (2.43 ± 0.08 vs. 2.44 m/s; One-sample t-test; t(38) = −0.30, p = 0.76). Conduction speed increased exponentially with the addition of each new sheath, and the largest gain was seen with the creation of a completely continuous stretch of myelin (**Fig. 5i,j; Supplementary Fig. 6**). Together, these results indicate that sheath retraction, resulting in increased node length, may slow or halt conduction along certain axons during the acquisition of a new motor task. In contrast, after motor learning, heightened generation of continuous myelination may speed up transmission.

Our experimental data suggests staged myelin dynamics can result in one of three outcomes in the two weeks following learning (**Fig. 2i, j**): 1) a decrease in overall myelin coverage due to node lengthening during learning; 2) the maintenance of overall myelin coverage due to node lengthening during and after learning and sheath addition after learning; and, 3) an overall increase in myelin coverage due to sheath addition following learning. In each of these three conditions, our modeling suggests that node lengthening will slow conduction during learning. After learning, former nodes continue to lengthen, and, if no new sheaths are added, conduction will continue to decrease. In contrast, if new sheaths are added to the axon, our modeling suggests that conduction increases. The magnitude of these changes in conduction speed depends on the number and distribution of ion channels along the lengthening gap in myelination. Higher leak conductance and lower sodium channel conductance will lead to conduction delays and, less often, failure to propagate (**Fig. 5**). Post-hoc analyses of lengthening nodes using pan-sodium channel isoform antibodies found examples with and without the presence of sodium channel clusters. We analyzed over 100 gaps in myelin of a similar distance in fixed tissue and found that only 52.9 ± 4.58% possessed distributions of sodium channels (**Supplementary Fig. 3d,e**). These results suggest that clusters of sodium channels may either be redistributed along newly lengthened gaps in myelin or that nodal complexes may disintegrate as the node lengthens. Together, our post-hoc analyses and modeling suggest that both gap length and distribution of sodium channels together can be finely modulated to produce highly specific changes to conduction during learning.

### Learning-activated axons show heightened myelin and nodal plasticity

Previous studies show that optogenetic and chemogenetic activation of neurons can drive oligodendrogenesis^4^ and myelination of specific axons^12^ in the adult brain. Yet, how myelination is modified by behavior on specific neuronal ensembles associated with learning or life experience remains unclear. To answer this question, we labeled learning-activated axons using an activity- and tamoxifen-dependent viral approach called Targeted Recombination in Active Populations (TRAP)^29,30^. We combined this approach with longitudinal, *in vivo* imaging to monitor the dynamic changes in myelin and nodes on activated neuronal ensembles involved in the forelimb reach. We injected L2/3 of the primary motor cortex of *MOBP-EGFP;R26-lsl-tdTomato* reporter mice with a cFos-ERT2-Cre-ERT2-PEST AAV, which drives tamoxifen-dependent Cre expression via the activity-dependent cFos promoter^31^. Longitudinal, *in vivo* imaging allowed us to identify neurons labeled by tamoxifen-independent recombination and separate these false-positives (baseline labeled; **Fig 6a,b**) from the learning-activated condition (learning activated; **Fig. 6a,b**). This approach revealed tamoxifen-independent tdTomato expression plateaued ~40 days following viral injection (**Supplementary Fig. 7**). Therefore, six weeks following viral injection, mice were trained on a forelimb reach task for seven days and 4-hydroxytamoxifen (4-OHT) or vehicle was delivered three hours after the final session of motor learning to label the learning-activated neuron ensemble associated with the forelimb reach, as previously described^31^. Importantly, we found that learning-activated neurons and their associated axons were permanently labeled with tdTomato only in mice that received both motor learning and 4-OHT administration. Untrained mice that received 4-OHT or trained mice that received vehicle did not show an increase in the number of neurons expressing tdTomato (**Supplementary Fig. 7**). The majority of learning-activated local neurons were found in Layer 2/3 of cortex with descending projections (likely IT-type pyramidal neurons^32^) (**Supplementary Fig. 7**). Using this approach, we traced the proximal 240±86 microns of the axon of learning-activated neurons to determine changes in myelination of neuronal ensembles associated with motor learning (**Fig. 6c; Supplementary Fig. 7**).

**Figure 6.**
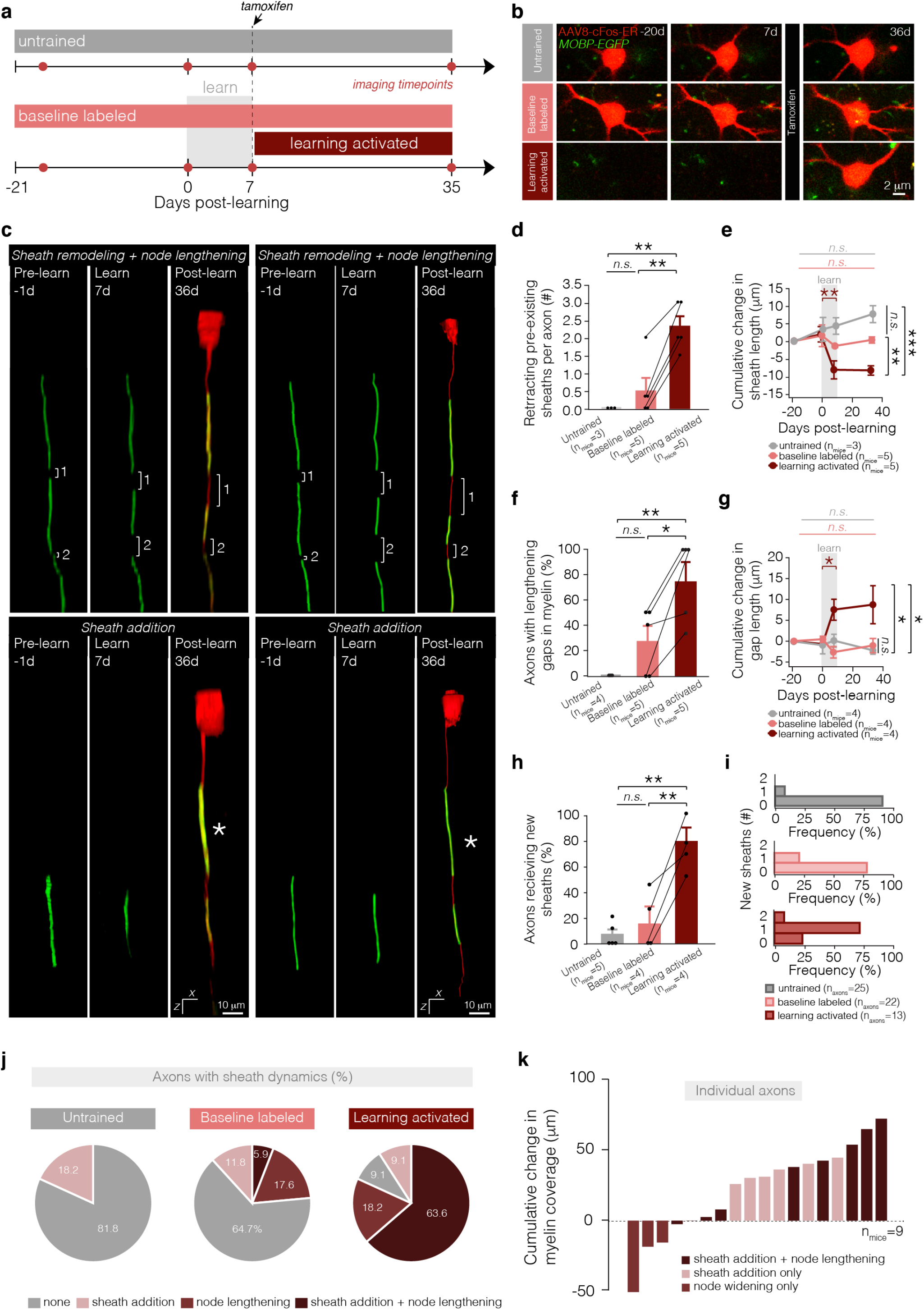
**c,** Reconstructions of Layer 2/3 axons exhibiting coupled node lengthening and sheath retraction (top) and sheath addition (bottom) following learning. See **Supp. Fig. 8** for raw data associated with reconstructions. **d,** Learning modulates number of retracting sheaths per axon (F_2,26.19_ = 6.65, p = 0.0007). One month after learning, learning-activated axons have significantly more sheath retraction relative to axons labeled at baseline in learning mice (p = 0.0007; Tukey’s HSD) and in untrained mice (p = 0.0062; Tukey’s HSD). Dot and error bars represent median and I.Q.R. **e,** Learning modulates dynamics of sheath retraction (F_6,7.14_ = 28.07, p < 0.0001). During learning, sheaths on behaviorally-activated axons retract significantly (p = 0.0065; Tukey’s HSD), and one month after learning, sheaths are significantly shorter than sheaths on baseline-labeled axons in learning mice (p < 0.0026; Tukey’s HSD) and in untrained mice (p < 0.0001; Tukey’s HSD). **f,** Learning modulates the proportion axons with lengthening nodes (F_2,14.68_ = 2.96, p = 0.029). One month after learning, a significantly higher proportion of learning-activated axons have node lengthening relative to axons labeled at baseline in learning mice (p = 0.030; Tukey’s HSD) and in untrained mice (p = 0.044; Tukey’s HSD). **g,** Learning modulates dynamics of node lengthening (F_6,4.76_ = 24.32, p = 0.0025). During learning, nodes on learning-activated axons retract significantly (p = 0.022; Tukey’s HSD), and one month after learning, nodes are significantly longer than nodes on baseline-labeled axons in learning mice (p = 0.047; Tukey’s HSD) and in untrained mice (p = 0.020; Tukey’s HSD). **h,** Learning modulates the proportion axons with new sheath addition (F_2,25.02_ = 8.70, p = 0.0002). One month after learning, a significantly higher proportion of learning-activated axons have new sheath addition relative to axons labeled at baseline in learning mice (p = 0.0006; Tukey’s HSD) and in untrained mice (p = 0.0004; Tukey’s HSD). **i, The majority of learning-activated axons receive 1 new sheath following training. j,** Proportion of axons showing no dynamics (grey), only sheath addition (pink), only node lengthening (red), or both sheath addition and node lengthening (dark red). **k,** Cumulative change in myelin coverage on individual axons from baseline to four weeks post-learning. *p < 0.05, **p < 0.01, ***p < 0.0001, NS, not significant; bars and error bars represent mean ± s.e.m unless otherwise noted. For detailed statistics, see **Supplementary Table 3, Figure 6**.

While the myelination profile did not differ between conditions in learning mice before motor training (75±14.4 vs. 70±13.3% of axons with intermittent myelination; Student’s t test, t(6.67) = 0.25, p = 0.81), we found that learning-activated L2/3 axons showed increased numbers of retracting pre-existing myelin sheaths following motor learning compared to baseline-labeled (tamoxifen-independent) tdTomato-positive neurons present at the onset of *in vivo* imaging in untrained mice and in mice that learned (**Fig. 6d,e**). This sheath retraction was temporally correlated to learning, resulting in a significant decrease in sheath length during the learning period (**Fig. 6e**). These sheath dynamics resulted in increased numbers of lengthening unmyelinated gaps on learning-activated, sparsely myelinated axons relative to baseline-labeled L2/3 axons in untrained mice and in mice that learned (**Fig. 6f**). Similar to sheath retraction, gap lengthening was also temporally correlated with learning, and on learning-activated axons, the gaps between myelin sheaths increased significantly, lengthening 8.32±2.38 microns during the learning period and achieving significantly larger lengths by one month following learning than unmyelinated gaps on baseline-labeled axons in untrained mice and mice that learned (**Fig. 6g**). Next, we examined changes in the addition of new myelin sheaths on learning-activated axons. Following learning, a higher proportion of learning-activated axons exhibited new sheath addition relative to baseline-labeled axons in untrained mice and learning mice (78.33±21.73 vs. 9.08±6.22 vs. 14.49±8.09% per mouse; **Fig. 6h,i**). We found that a large proportion of learning-activated axons (63.6%) show both sheath addition, and node lengthening on the same axonal segments compared to zero axons in untrained mice and 5.9% axonal segments in baseline-labeled axonal segments in learning mice (**Fig. 6j**). These data indicate that myelin and nodal plasticity is occurring on the same segments of learning-activated axons in trained mice. These myelin dynamics often resulted in an overall increase in myelin coverage relative to baseline (6/9 axons with node lengthening and sheath addition; **Fig. 6k**). These results suggest that altered sheath placement, rather than heightened oligodendrogenesis, drives increased sheath addition on learning-activated axons in Layer 2/3 of primary motor cortex following learning. Taken together, these data suggest that the diversity in patterns of intermittent myelination on cortical axons may be explained in part by activity-dependent recruitment of myelin sheaths and plasticity of pre-existing myelin and nodal distributions.

### The magnitude and timing of myelin dynamics correlate with behavioral performance

During training of the dexterous reach task, the success rate (**Supplementary Fig. 2**) and degree of myelin dynamics (**Fig. 2i**) can vary over the course of training between individual mice. To explore whether the extent of myelin plasticity is linked to behavioral performance, we compared the magnitude of node lengthening to the number of days above 20% success in the reach task during learning for individual mice (**Fig. 7a**). We find that node lengthening is strongly correlated with reach task proficiency. Similarly, myelin sheath addition is strongly correlated with behavioral proficiency two weeks following learning (**Fig. 7b**). Next, we explored whether a staged myelin response (nodal plasticity followed by sheath addition, **Figs. 2,4**) may be characteristic of high but not low behavioral performance. To do this, we analyzed the temporal dynamics of node lengthening and sheath addition on the population of axons that showed changes to myelin in response to learning. We classified animals according to behavioral performance on the reach task: animals which attained 3 or more days above 20% successful reach attempts were considered to have high performance. In these mice, sheath addition was significantly delayed relative to node lengthening (**Fig. 7c**), where the majority of node lengthening occurred during learning (half maximum: 6.42 ± 1.2 days following onset of learning) and the majority of sheath addition began in the two weeks following learning (half maximum: 12.00 ± 1.09 days following onset of learning). In contrast, in animals with low performance (fewer than 3 days above 20% successful reach attempts), there was no staged response of node lengthening followed by sheath addition (**Fig. 7d**). In mice with low performance, both processes proceeded linearly and were not shifted in time relative to one another (half maximum: 9.68±1.97 vs. 9.20±0.881 days following onset of learning for node lengthening and sheath addition, respectively). Together, these results indicate a strong relationship between both the magnitude and timing of myelin and nodal dynamics in forelimb motor cortex and behavioral performance on a fine-skilled motor task.

**Figure 7.**
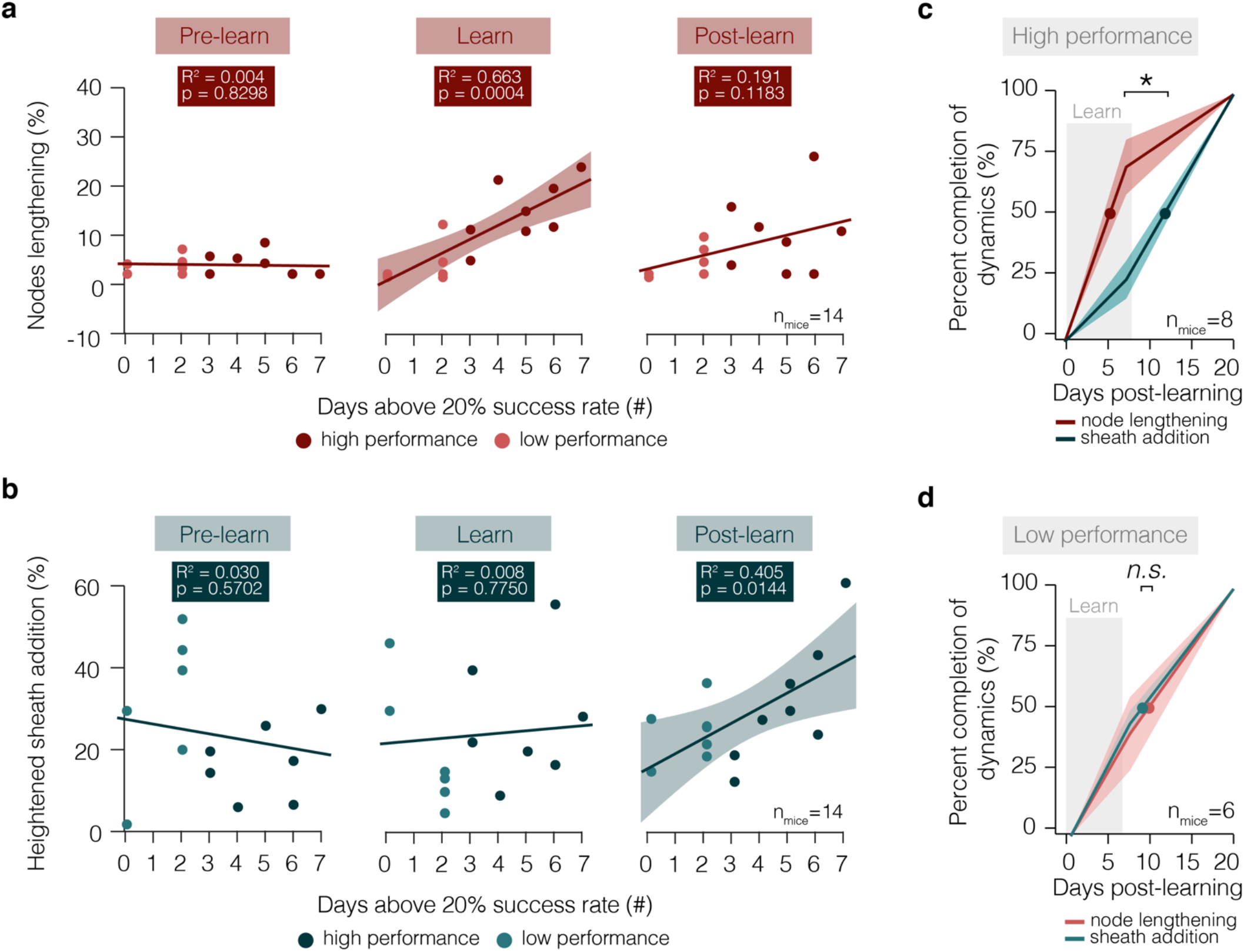
The extent and timing of myelin dynamics is correlated to behavioral performance. **a,** The proportion of nodes lengthening per mouse is correlated to degree of behavioral success only during learning, but not before or in the two weeks after learning. **b,** The proportion of axons receiving new myelin sheaths per mouse is correlated to degree of behavioral success only in the two weeks following learning, but not before or during learning. **c,** In high performing animals (defined as animals which attain 3 or more days above 20% successful reaches), node lengthening occurs significantly earlier than sheath addition (Paired Student’s t-test; t(5)=3.99, p=0.11). **d,** In low performing animals (defined as animals which attain less than 3 days at above 20% successful reaches), node lengthening and sheath addition occur concurrently. *p < 0.05, **p < 0.01, ***p < 0.0001, NS, not significant; lines and shaded areas represent mean ± s.e.m; dots represent half maximum. For detailed statistics, see **Supplementary Table 3, Figure 7**.

## Discussion

Changes to nodal and myelin sheath properties are emerging as potential regulators of conduction, suggesting that experience-driven changes to patterns of myelination may play a role in shaping neuronal communication during learning. Exploring learning-induced myelin dynamics on learning-relevant axons is vital to understanding how changes to myelin might drive circuit dynamics during learning. Here, we establish that two forms of learning-related myelin plasticity— nodal dynamics and new sheath addition—are specifically targeted to motor cortical axons activated by motor learning in the healthy adult mouse. Learning-induced myelin plasticity proceeds in a staged response: during learning, pre-existing sheath retraction creates new patterns of intermittent myelination on activated axons, while new sheaths from oligodendrocytes generated after learning preferentially eliminate large gaps in pre-existing myelin. Furthermore, we provide a biologically-relevant computational model which suggests that plastic changes to myelin pattern, nodal distribution, and ion channel properties along unmyelinated gaps may have profound effects on action potential propagation and circuit function. Lastly, we define a novel relationship between behavioral performance and the timing and magnitude of myelin and nodal plasticity.

Remodeling of pre-existing myelin sheaths was first proposed as a mechanism underlying the observation that adult-born oligodendrocytes generated shorter myelin segments than oligodendrocytes generated during development in the highly myelinated optic nerve^33^. Subsequent longitudinal, *in vivo* imaging studies confirmed that pre-existing myelin sheaths could be remodeled and documented the dynamics of this process in the context of sensory deprivation or aging^5,22,34^. Yet, the link between changes in neuronal activity and the remodeling of pre-existing myelin sheaths remains unclear. Our studies indicate that myelin remodeling occurs specifically on myelinated axons with learning-induced neuronal activity. These findings support previous work showing that generalized modifications to neuronal activity, such as changing sensory experience via monocular deprivation, modulate pre-existing myelin sheath length in a neuronal subclass-dependent manner^22^. During the formation of new myelin sheaths following oligodendrocyte generation, axonal vesicular release is key in regulating the growth of nascent sheaths^10,11^. This is controlled in part by modulating local calcium concentration within individual new myelin sheaths^13,14^. However, synaptic vesicle release regulates the growth of myelin sheaths on certain glutamatergic axons but not others^35^ suggesting neuron-specific or additional oligodendrocyte-intrinsic cues may play important roles in sheath formation. Indeed, oligodendrocyte-intrinsic cues direct regional differences in myelin sheath lengths of cortical-versus spinal cord-derived oligodendrocytes^36^. These cues may include ion channel signaling via hyperpolarization-activated, cyclic nucleotide-gated (HCN) ion channels, and inhibitory transmembrane proteins such as Nogo-A, which modulate length of new myelin sheaths^37,38^. Future work investigating whether similar signaling mechanisms regulate changes in the length of pre-existing myelin sheaths on learning-activated axons will provide additional insight into the fundamental biology regulating myelin remodeling.

Building evidence suggests that the patterns of myelination that emerge during development may be used to fine tune conduction properties of specific axons^16^. Furthermore, the varied distribution of lengths of nodes of Ranvier in the optic nerve and corpus callosum suggests that dynamically altering nodal parameters will alter conduction velocity^17^. Our data indicate that motor learning induces changes in node length, mediated through sheath retraction, specifically on the axons of learning-activated neurons. How these dynamic changes modify ion channel composition of nodes of Ranvier or heminodes remains unclear. In the case of myelin removal through demyelination, the distribution and composition of ion channels are destabilized leading to changes in axonal excitability (reviewed by Waxman^39^). At learning-induced lengthened nodes, we find a broad range of distributions of sodium channel clusters at dynamic nodes (**Supplementary Fig. 3**). Across a range of possible sodium and leak conductances, our computational modeling predicts that retraction of pre-existing myelin sheaths will alter conduction speed, suggesting that dynamic changes in myelination patterns may play an important role in cortical circuit function. Furthermore, since assemblage of sodium and potassium channels localized to nodes varies depending on brain region, neuronal subclass, maturation, and myelination (reviewed by Suminaite et al.^40^), future work should consider these additional variables when exploring how learning-induced nodal dynamics alter axonal excitability and conduction.

The functional ramifications of learning-induced changes to myelination warrant further exploration. One possibility is that preexisting sheath remodeling during learning may serve to introduce variability in early stages of learning^41^. For example, altering spike-timing plasticity via changes in action potential conduction could modify circuit function. Although LTD and LTP can occur with prepost spike delays of up to ±100 millisecond (ms), spike timingdependent plasticity operates most strongly within a 10 ms window, with a sudden 1-5 ms transition between LTP and LTD^42^. This suggests that even small shifts in arrival time—as predicted with alterations to node length or sheath addition—could be significant, changing not only the strength of the synaptic connection but also potentially determining whether a synapse is strengthened or weakened (see Fields^43^). Indeed, in the songbird forebrain minor delays in conduction can influence the sequential activation of neuronal ensembles to profoundly shape communication within a circuit^44^. Here, we show that retraction of continuous myelin sheaths results in the generation of new areas of intermittent myelination during learning, and our modeling suggests this slows conduction during task acquisition. After learning, we find elimination of pre-existing gaps in myelin by new sheaths to generate stretches of continuous myelination, resulting in an increase in conduction speed. Interestingly, the deposition of new myelin increases axon caliber during development in the peripheral nervous system^45,46^, potentially serving as an additional mechanism to modulate conduction velocity. Together, our data indicate that the transition from partial to full myelination, and vice versa, may be important to dynamically alter neuronal communication when acquiring a new motor skill.

We previously reported that the generation of new oligodendrocytes in the forelimb motor cortex increases following the completion of reach task training^20^. Consistent with these results, here we found that new sheath addition increases after attaining proficiency in the reach task and correlates with behavioral performance suggesting this process may perform a distinct role in the late phases of learning. Indeed, recent work indicates that the generation of new oligodendrocytes regulates memory consolidation^8,9^. In addition, we replicate and extend previous results indicating that oligodendrogenesis is heightened in L1 but not L2/3 of motor cortex in response to learning^20^. Our current data show that the proportion of L2/3 axons receiving new myelin sheaths was increased following learning indicating that the targeted placement of new sheaths is independent of the rate new oligodendrocyte formation. Together, these data suggest regional differences in learning-related myelin plasticity may reflect varying needs depending on cortical layers and circuits.

Our computational modeling suggests new sheath addition may serve to speed up conduction along consolidated neural circuits after learning. However, it is important to consider that myelin serves additional roles beyond influencing conduction speed, including metabolic support of axons^47,48^. Recent studies suggest that oligodendroglial metabolic support of axons contributes to information processing^49^. The high density of connections in the cortex results in substantial energetic demands^50^ and activity-dependent myelin sheath addition may reflect increased metabolic needs due to heightened neuronal activity. Previous studies show that lactate, an oligodendrocyte-derived source of metabolic support to axons^47,48,51^, can rescue functional decline in the optic nerve due to glycogen depletion^52^. Furthermore, since myelin acts as a barrier to accessing extracellular energy supplies^53,54^, removal of myelin via sheath retraction may be driven by metabolic need, exposing axonal membrane to allow access to extracellular glucose. Exploring the effects of learning-induced myelin and nodal dynamics in the context of neuronal metabolic demands will improve our understanding of the roles of oligodendrocytes and myelination in information processing.

Another possible role for learning-induced nodal and myelin remodeling is to regulate the plasticity of axon morphology, signaling, and synaptic connections. On PV interneurons, the degree of myelination is correlated to both branch morphology and the number of *en passante* boutons^55^. Furthermore, on intermittently myelinated L2/3 pyramidal neurons, unmyelinated regions contain both afferent and efferent synapses^2^. The staged response of learning-related myelin dynamics, where the retraction of existing myelin and suppression of oligodendrogenesis occurs during learning^20^, may enable the formation of new synaptic connections in motor cortex or local initiation or modulation of action potentials^56,57^. Subsequently, targeted sheath addition following learning could lead to the closure of a learning-related window of heightened plasticity, as suggested in studies modulating the duration of sensory map plasticity using Nogo receptor (NgR) knockout mice^58^. This staged plasticity may occur in an axon-specific manner as we find a synchronization of nodal dynamics along specific axonal segments as well as targeted new myelin sheath addition. These findings suggest that individual axon segments have heightened levels of myelin plasticity similar to location-dependent synaptic plasticity on dendritic branches which display learning-induced branch-specific calcium spikes and clustered dendritic spine formation^59,60^. While chemogenetic stimulation of parvalbumin-positive axons drives newly-generated axonal collaterals that receive *de novo* myelination^61^, recent work shows pre-existing myelin dynamics are independent of increased axon branching events in visual cortex following monocular deprivation^22^. Additional information examining the specific timing of myelin remodeling and neuronal morphological plasticity will aid in determining direct and indirect effects of these processes.

Exploring the link between learning-relevant neuronal activity and changes in myelination has important implications for myelin repair in the context of neurological disease. In multiple sclerosis, disability is lessened as remyelination increases^62^, but the extent of remyelination of lesions can differ within the same individual^63,64^. Furthermore, following acute demyelination, the pattern of myelination is often re-established in regions of continuous myelination, while regions of discontinuous myelination are less likely to be replaced^65,66^. Together, these data suggest that lesions comprising different neuronal circuits may experience different levels of myelin repair. Here, we show that new myelin sheath addition and changes to pre-existing myelin sheath length are targeted to axons in a learning-activated circuit. Our work establishes that behavior shapes the pattern of myelination on the level of individual axons within defined, activated circuits in the healthy brain. These findings raise the possibility that learning-induced neuronal activity may modulate the specificity and efficacy of myelin repair following a demyelinating injury. Understanding how and when the brain engages in plastic changes to myelin could inform interventions to address myelin-altering pathology, as seen in multiple sclerosis and stroke, where targeted myelin dynamics hold potential to improve therapeutic success.

## Acknowledgements

We thank Mike Hall for machining expertise; Helena Barr for the conversation that re-initiated interest in labeling activated axons; and past and current members of the Hughes and Welle labs and the CU Anschutz Myelin Group for discussions.

## Author Contributions

EGH and CMB conceived the project. CMB designed and performed experiments, analyzed data, and generated all figures. RH performed all viral injections and performed surgeries. LAO performed experiments and analyzed data for Fig. 7, Supplementary Fig. 3. MAT performed surgeries. LC performed experiments and additional imaging analyses for Figs. 4,6,7, Supplementary Fig. 3. AC provided technical support for all mouse lines. APP supervised the computational modulating and EGH supervised the project. CMB and EGH wrote the manuscript with input from other authors.

## Data and materials availability

All data that support the findings, tools and reagents will be shared on an unrestricted basis; requests should be directed to the corresponding author.

## List of Supplementary Materials

Supplementary Tables 1 to 3

## Methods

### Animals

All animal experiments were conducted in accordance with protocols approved by the Animal Care and Use Committee at the University of Colorado Anschutz Medical Campus. Male and female mice used in these experiments were kept on a 14h light/10h dark schedule with ad libitum access to water and food, except during training-related food restriction (see *Forelimb Reach Training).* Mice were randomly assigned to experimental groups and were precisely age-matched across groups. *R26-lsl-tdTomato* reporter mice (Ai9; JAX: 007909), congenic C57BL/6N *MOBP-EGFP(MGI:4847238)* lines, which have been previously described^5,20^, were used for two-photon imaging.

### Two-photon microscopy

Cranial windows were prepared as previously described^67^. Induction of anesthesia in six-to eight-week-old mice was accomplished with isoflurane inhalation (induction, 5%; maintenance, 1.5–2.0%, mixed with 0.5L/min O2), and mice were kept at 37 °C body temperature with a thermostat-controlled heating plate. The skin over the right cerebral hemisphere was removed, the skull was cleaned, and a 2 × 2 mm region of skull centered over the forelimb region of the primary motor cortex (0 to 2 mm anterior to bregma and 0.5 to 2.5 mm lateral) was opened using a high-speed dental drill. A piece of cover glass (VWR, No. 1) was then placed in the craniotomy and sealed with Vetbond (3M) followed by dental cement (C&B Metabond). Carprofen (5mg/kg) was administered subcutaneously prior to awakening and for three additional days for analgesia. To stabilize the head during imaging, a custom metal plate with a central hole was attached to the skull. *In vivo* imaging sessions began 2-3 weeks post-surgery and took place once per week. At each imaging time point, mice were anesthetized with isoflurane and immobilized by attaching the head plate to a custom stage. Images were collected using a Zeiss LSM 7MP microscope equipped with a BiG GaAsP detector using a mode-locked Ti:sapphire laser (Coherent Ultra) tuned to 920 nm. The average power at the sample during imaging was 5-30 mW. Vascular and cellular landmarks were used to identify the same cortical area over longitudinal imaging sessions. Image stacks were acquired with a Zeiss W “Plan-Apochromat” 20X/1.0 NA water immersion objective giving a volume of 425 μm × 425 μm × 336 μm (2,048 × 2,048 pixels, 1.5 μm step size; corresponding to layers I-III, 0-336 μm from the meninges, or layer 2/3 110-446 μm from the meninges).

### Forelimb reach training

Mice were weighed, habituated to a training box for 20 minutes, and deprived of food 24 hours prior to training. The training box was fitted with a slot providing access to a pellet located on a shelf 1cm anterior and 1mm lateral to the right-hand side of the slot. After one session of initial habituation, training sessions began daily for 20 minutes or 20 successful reaches, whoever occurred first. Mice learned to reach for the pellet using their left hand. Successes were counted when the mouse successfully grabbed the pellet and transported it inside the box. Errors were qualified in three ways: “Reach error” (the mouse extends its paw out the window but does not grab the pellet), “Grasp error” (the mouse reaches the pellet but does not successfully grasp onto it), and “Retrieval error” (the mouse grasps the pellet but does not succeed in returning it to the box). Mice were kept on a restricted diet throughout training to maintain food motivation but were weighed daily to ensure weight loss did not exceed 10%. For forelimb reach training, mice underwent habituation (average of ~1 days of exposure) followed by training until seven consecutive days of training with reach attempts were recorded. Similar to previously published findings, over 90% of mice trained in forelimb reach context were able to learn the task^68^; mice were excluded if they were not able on a single day to succeed in at least 10% of reaches (n = 2 mice; **Supplementary Fig. 2**). To control for any experimenter effects in forelimb reach training results, all mice were trained by the same experimenter (i.e. control and experimental mice were only compared if trained by the same experimenter).

### Labeling of learning-activated axons

*R26-lsl-tdTomato* or *MOBP-EGFPR26-lsl-tdTomato* reporter mice were injected with 1ul of AAV-8-cFos-ERT2-Cre-ERT2-PEST^31^ in which CreER expression is driven by the activity-dependent *Arc* promoter at a 1:8 dilution (stock virus titer: 8.8 x 10^12^). 6 weeks later, mice were trained in a forelimb reach task (see *Forelimb reach training*). Age-matched controls were exposed to the box, fasted, and injected with tamoxifen but not given reach training. 10 mg/kg of 4-OHT was given to all mice three hours after the last training session to enable CreER-mediated recombination. Mice were then imaged for an additional 4 weeks following tamoxifen to allow the full expression of tdTomato fluorophore (see **Supplementary Fig. 7**).

### Near-infrared branding

Near-infrared branding (NIRB) marks were generated using our two-photon microscope (described above in *Two-photon microscopy*). The orientation and size of NIRB marks was regulated by modulating laser power and exposure time, as described previously (Bishop et al., 2011). To generate NIRB marks, we used a Zeiss W “Plan-Apochromat” 20X/1.0 NA water immersion objective. To locate cells and structures of interest within the longitudinally-imaged window, we burned a ~500 μm × 500 μm box with asymmetrical markings, ensuring burn marks were 100 μm away from the area of interest (**Supplementary Figs. 1,3**). We then captured a low-resolution two-photon image stack of the completed NIRB marks and their relationship to longitudinally-imaged structures to locate them across *in vivo* images and in post-hoc immunostained sections, using emission filters for yellow-green fluorescent proteins and DsRed. Prior to marking the brain with NIRB, mice were anesthetized with an intraperitoneal injection of sodium pentobarbital (100mg/kg b.w.) and transcardially perfused with 4% paraformaldehyde in 0.1 M phosphate buffer (pH 7.0-7.4).

### Correlated *in vivo* and immunohistochemistry

Brains were postfixed in 4% PFA for 1 hr at 4 °C, transferred to 30% sucrose solution in PBS (pH 7.4), and stored at 4 °C for at least 24 h. Brains were extracted, frozen in TissuePlus O.C.T, and sectioned at 30 μm thick. To obtain correlated images of immuno-processed sections, we first micro-dissected out the cranial window area and oriented the tissue to be cryosectioned in a plane parallel to the surface of the imaged area of the brain. Once sectioned, we selected fixed cryosections to immunostain by locating the NIRB markings on a confocal microscope (Zeiss LSM 510).

Immunostaining was performed on free-floating sections. Sections were pre-incubated in blocking solution (5% normal donkey serum, 2% bovine g-globulin, 0.3% Triton X-100 in PBS, pH 7.4) for 1-4 h at room temperature, then incubated overnight at 4 °C in primary antibody (listed along with secondary antibodies in **Supplementary Table 2**). Secondary antibody incubation was performed at room temperature for 2 h. Sections were mounted on slides with Vectashield antifade reagent (Vector Laboratories). Images were acquired with a laser-scanning confocal microscope (Zeiss LSM 510).

### Image processing and analysis

Image stacks and time-series were analyzed using FIJI/ImageJ. All analysis was performed on single plane images, which were pre-processed by with a Gaussian blur filter (radius = 1 pixel) to denoise and aid identification of individual myelin sheaths and nodes. When generating figures, image brightness and contrast levels were adjusted and maximum projections were used for clarity. Longitudinal image stacks were registered using FIJI plugins ‘Correct 3D drift’^67^ or ‘PoorMan3DReg’. When possible, blinding to experimental condition was used in analyzing image stacks from two-photon imaging.

### Myelin sheath and node analysis and analysis of myelin profiles along single axons

*In vivo* z-stacks were collected from *MOBP-EGFP* mice using two-photon microscopy. Z-stacks were processed with a 1-pixel Gaussian blur filter to aid in the identification of myelin internodes. Myelin paranodes and nodes of Ranvier were identified as described previously^5^, by increase in fluorescence intensity for paranodes and a decrease to zero in EGFP fluorescence intensity for nodes. Node length was defined as the distance between each paranodes as measured by Simple Neurite Tracer (see **Supplementary Fig. 3**). To identify all sheaths belonging to a single oligodendrocyte, processes connecting the oligodendrocyte cell body to each individual sheath were traced, as previously described^20^. In *MOBP-EGFP* mice, axon morphology was determined using the trajectory of the continuously myelinated axon and these traces were validated in a subset of mice using post-hoc immunostaining for axons in the same tissue (see **Supplementary Fig. 3**). Two-color images of tdTom+ axons and their associated myelin sheaths (labeled with EGFP) were registered using FIJI plugins ‘Correct 3D drift’ or ‘PoorMan3DReg’. Axons, myelin, and nodes were then traced using Simple Neurite Tracer at baseline, the first day of training, the last day of training, and either 2 or 4 weeks following training (see **Supplementary Fig. 7**). Neurons, oligodendrocytes, and sheaths that resided within a volume of425 × 425 × 110 μm^3^ from the pial surface were considered in Layer 1 and those at a depth of 110-446 μm^2^ were considered Layer 2/3. Sheaths were defined as new or pre-existing. Persisting sheaths and nodes lasted for the entire imaging time course; new sheaths appeared after the onset of imaging. Superficial motor cortex is sparsely myelinated, and, at the resolution of our imaging volume, very few sheaths occupied completely overlapping territories. However, in the rare event in which a sheath’s trajectory could not be determined, it was excluded from analysis.

### Computational modeling

We used the NEURON simulation environment (v.7.4; Hines & Carnevale, 1997) to simulate action potential propagation along a myelinated axon. Details of the parameters used are summarized in **Supplementary Table 1**. The axon is divided into compartments representing the node and internode. 52 nodes and 51 myelin internodes were simulated, and conduction speed was measured between the 30^th^ and 40^th^ node.

Spatial resolution was set using 5 segments per cellular compartment, which ranged from 1-184 microns in length. Parameter bounds are described in **Supplementary Table 1**. Briefly, the lower and upper bounds for sodium conductance and leak conductance were defined using biologically relevant values consistent with previous publications^28^. Myelinated axon and node diameters were set to 0.902 microns and 0.72 microns, respectively, and were within the range of experimentally-derived values for myelinated cortical axons from previous publications^17,65,69^. Pre-learning (1 micron), learning (20 microns), and post-learning (35 microns) node lengths were set using experimentally-measured values (see **Figs. 1–3; Supplementary Fig. 6**). Similarly, myelin sheath length was set to 60 microns using experimentally-derived values (see **Supplementary Fig. 6**). A g-ratio of 0.7 was used, consistent with values determined from previous publications^65,69^. Temperature was set to 37° C, as in previous publications^17,70^. Hodgkin-Huxley style kinetics^71,72^ for NaV1.6 channels were fit using data from Huguenard et al.^73^ and Hamill et al.^74^ and Kv1 channel kinetics were fit based on rates from Hamill et al.^74^. Model files and equations were obtained from Hu et al.^70^.

### Statistics

A detailed and complete report of all statistics used, including definitions of individual measures, summary values, sample sizes, and all test results can be found in **Supplementary Table 3**. Sample sizes were not predetermined using statistical methods, but are comparable to relevant publications. All data were initially screened for outliers using IQR methods. All mice in a litter underwent cranial window surgery and concurrent two-photon imaging and training timelines were designated to be a “batch”. When possible, experimental groups were replicated in multiple batches with multiple experimental groups per batch. Statistical analyses were conducted using JMP (SAS). We first assessed normality in all datasets using the Shapiro-Wilk test. When normality was violated, we used nonparametric rank-sum tests. When normality was satisfied, we used parametric statistics including paired or unpaired two-tailed Student’s t-tests (depending on within- or between-subjects nature of analysis), or one-way and two-way ANOVA with Tukey’s HSD post-hoc tests. Two-tailed tests and α ≤ 0.05 were always employed unless otherwise specified. For statistical mixed modelling, we used a restricted maximum likelihood (REML) approach with unbounded variance component, and least-squared Tukey’s HSD post-hoc tests. All models were conducted with either one or two fixed effects, in which case we ran full factorial models. For all models, we used “Mouse ID” as a random variable unless otherwise specified. To test the significance of our regression models, we calculated the R-squared and F-test. Where we found significant effects, we subsequently calculated effect size using Cohen’s d, and these results may be found in **Supplementary Table 3**. For data visualization, all error bars represent the standard error of the mean unless otherwise specified.

## SUPPLEMENTARY MATERIALS

**Supplementary Figure 1.**
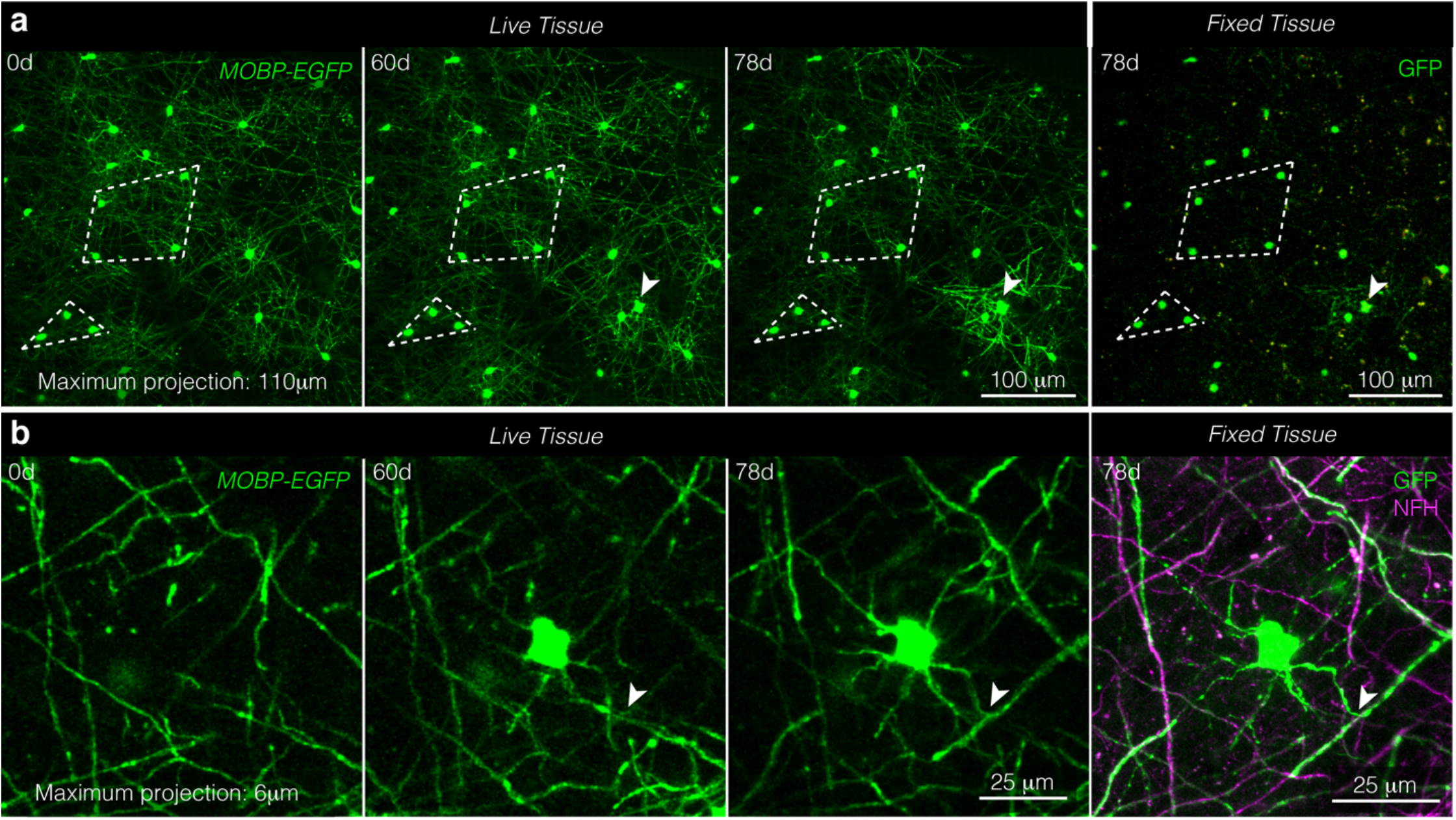
Near-infrared branding identifies the same oligodendrocytes and myelin sheaths in longitudinally, *in vivo* imaged areas and post-hoc stained tissue. **a,** The same field of view imaged *in vivo* (“Live Tissue”, left) and fixed, sectioned, and stained tissue (“Fixed Tissue”, right). Patterns of cell bodies (examples outlined in white dotted lines) were maintained across live and processed tissue. Note the new oligodendrocyte generated at 60d and delineated with a white arrowhead. **b,** A newly generated oligodendrocyte *in vivo* (“Live Tissue”, left) and fixed, sectioned tissue (“Fixed Tissue”, right) stained for oligodendrocytes and myelin (GFP, green) and axons (NFH, magenta). Note the same T-junction across live and fixed samples is marked with the white arrowhead.

**Supplementary Figure 2.**
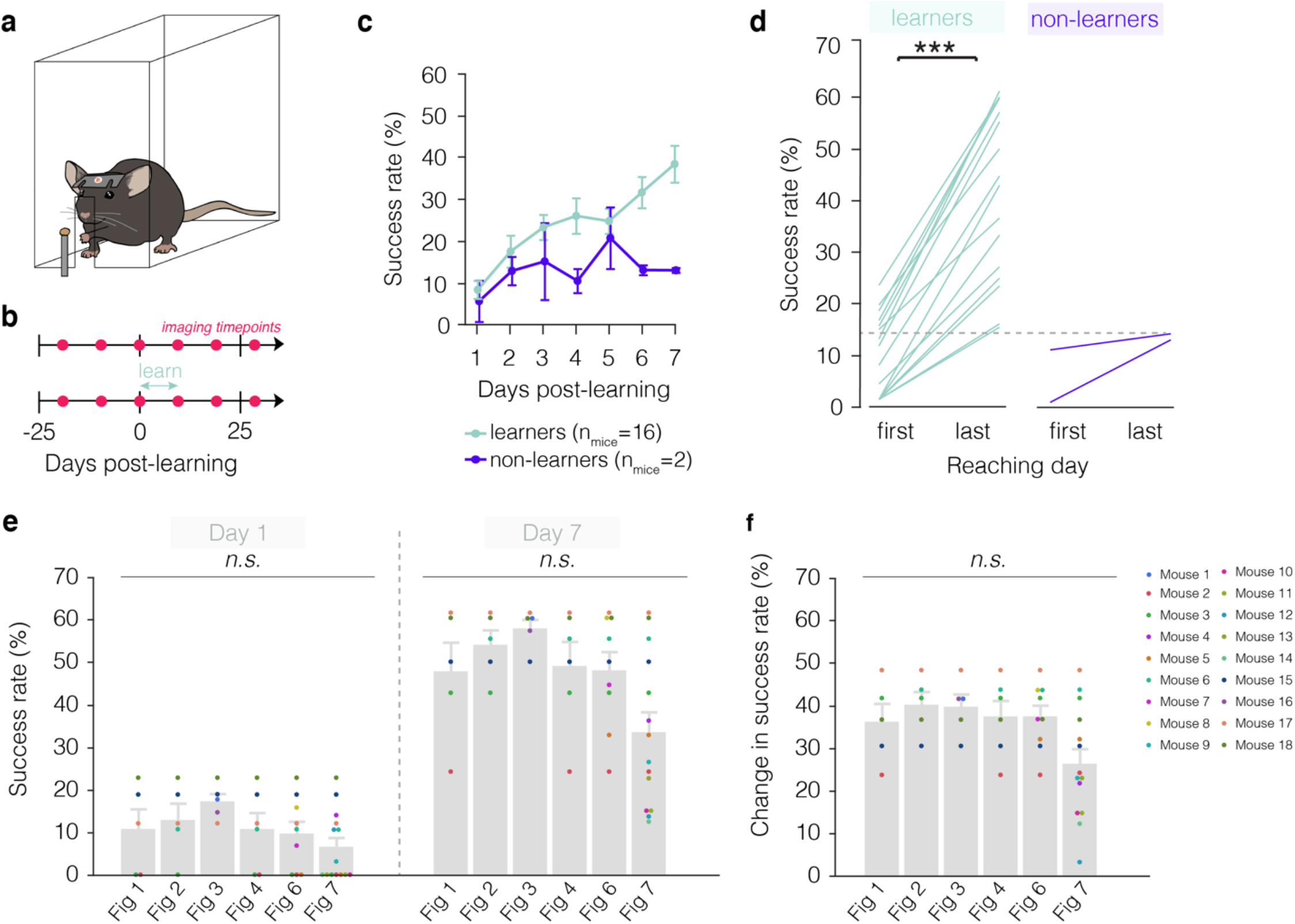
Learning trajectory of mice engaging in forelimb reach training. **a,** Illustration of forelimb reach training box environment. Mice learn to reach by extending their left hand through a slit in a plexiglass box to grab a pellet and return it to their mouth. **b,** Imaging and training timelines for untrained (top) and learning (bottom) mice. **c,** The majority of mice learn to perform the forelimb reach task (learners, green). Learners improve their success rate gradually over the course of seven days of training. In contrast, non-learners maintain a low success rate across time and do not attain higher than a 15% success rate at the end of the training regimen (purple). **d,** Successful learners of the task perform significantly better on the last day of training (Paired Student’s t-test, t(15)=11.72, p<0.0001) and achieve higher than 15% success rate, in contrast to non-learners which do not improve significantly and do not attain above 15% success on the last day of training. Mice were excluded from data analysis if they did not succeed in at least 10% of reaches across the seven days of training (n = 2 mice). **e, f,** Breakdown of all mice included in each of the figures, where each dot is a unique color that represents a single mouse and is consistent across graphs. Neither success rate (**e**) nor change in success rate (**f**) differ across figures.

**Supplementary Figure 3.**
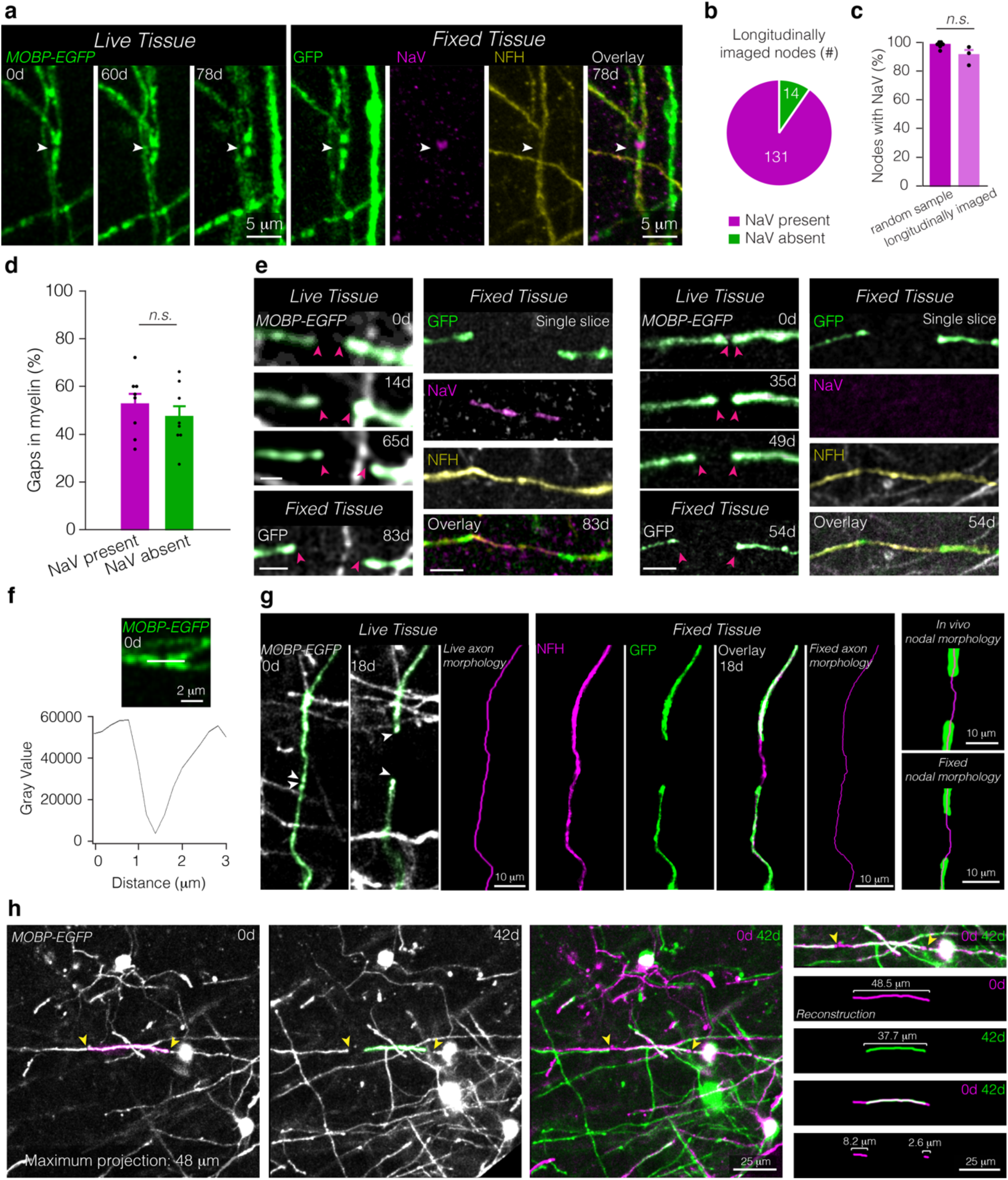
Identifying and tracing nodes and axons across live and fixed tissue. **a,** The same node in live imaged (“Live Tissue”, left) and fixed, sectioned tissue (“Fixed Tissue”, right) stained for oligodendrocytes and myelin (GFP, green), sodium channels (NaV, magenta), and axons (NFH, yellow). White arrow points to node across time and in fixed tissue. **b,c,** 90.90±3.73% of nodes visualized *in vivo* possess NaV staining characteristic of nodes of Ranvier (**b**), comparable to values of nodes identified fixed tissue of age-matched controls (**c**). **d,** No difference in the proportion of gaps in myelination (larger than 3 microns) with and without appreciable sodium channel distributions in fixed tissue of age-matched controls. **e,** Representative *in vivo* imaging and post-hoc immunostaining of a lengthening node with sodium channels (left) and without appreciable sodium channels (right), **f,** The same node confirmed with NaV staining in (a) is identified in live imaging by a marked decrease in autofluorescence between two GFP-labeled myelin paranodes. **g,** Axon morphology is accurately identified *in vivo* as confirmed by post-hoc immunostaining of the same tissue. A lengthening node *in vivo* (“Live Tissue”, left) and fixed, sectioned tissue (“Fixed Tissue”, middle). Axon trajectory is identified using the shape of the myelin sheath prior to node lengthening (“Live axon morphology”, magenta) and confirmed using post-hoc immunostaining of the same sheaths for axons (NFH, magenta) and myelin (GFP, green) and reconstructions of the axons (“Fixed axon morphology”, magenta). *In vivo* node morphology (“*In vivo* nodal morphology”, top right) is reconstructed and confirmed using post-hoc immunostaining and morphological reconstruction (“Fixed nodal morphology”, bottom right). **h,** Sheath length and change in sheath length is determined by aligning fields of view using fiduciary marks (e.g. cell soma) which extend across the entire duration of the study, including pre-existing oligodendrocytes, which maintain their position throughout the course of imaging (left). Change in sheath length is mirrored by a change in paranode position (right). To determine the change in sheath length, the distance between the initial and final paranodal position is traced using Simple Neurite Tracer and overlayed stacks.

**Supplementary Figure 4.**
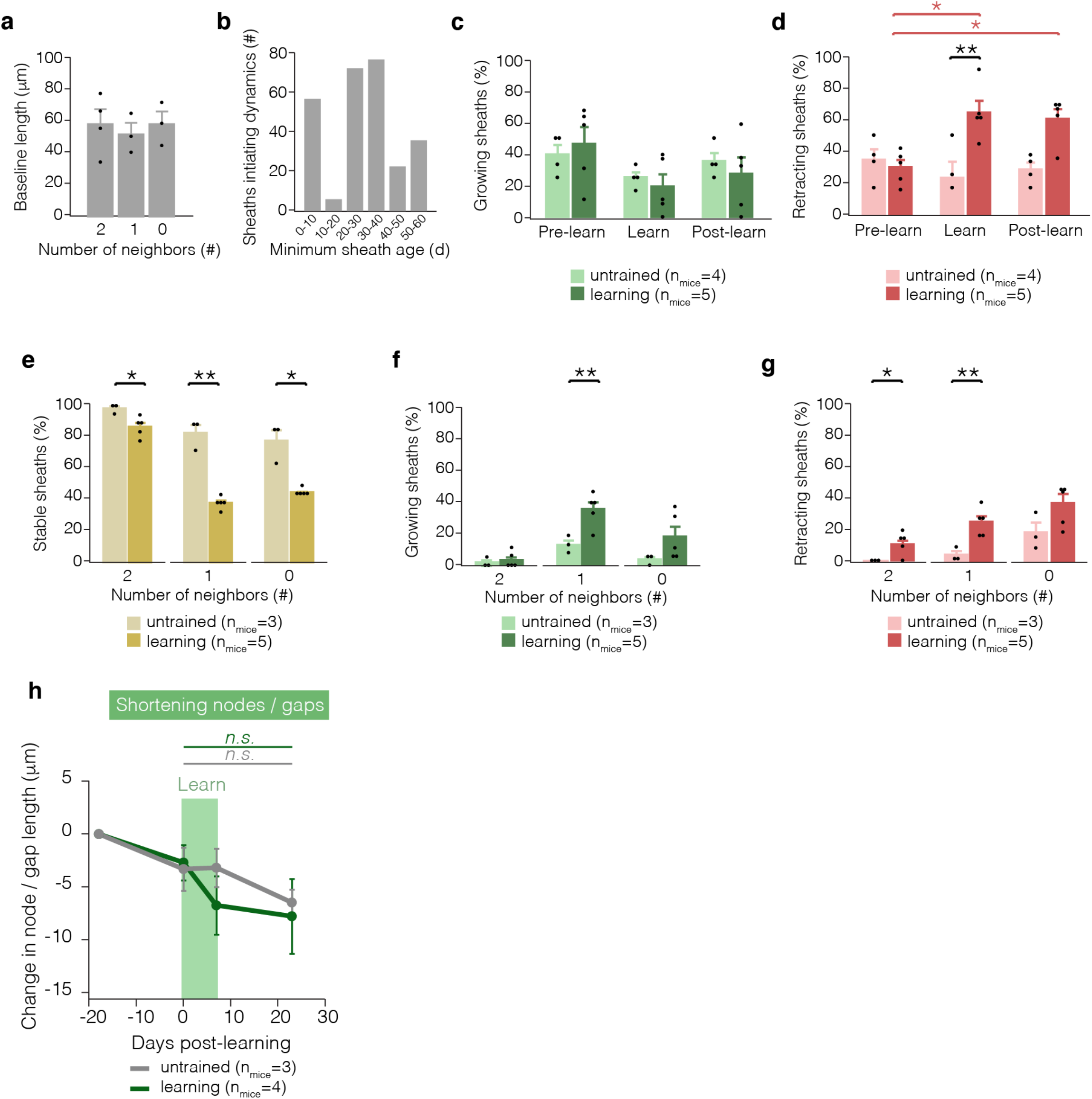
Sheath retraction, but not sheath growth, is affected by learning a new skill. **a,** Sheath length is similar across sheaths with 2, 1, or 0 neighbors. **b,** Sheaths of many ages initiate sheath dynamics in young adult mice. **c,** Proportion of dynamic sheaths engaging in growth three weeks before learning, during learning (one week), and in the two weeks after learning. **d,** Proportion of dynamic sheaths engaging in retraction before, during and after learning. Learning modulates sheath retraction (F_2,14_ = 6.76, p = 0.0088). During learning, more sheaths retract relative to untrained mice (p = 0.0095; Tukey’s HSD) and relative to pre-learning values in trained mice (p = 0.016; Tukey’s HSD). Two weeks after learning, more sheaths retract relative to pre-learning values in trained mice (p = 0.0358; Tukey’s HSD). **e,** In learning mice, there are fewer stable sheaths with 2 (p=0.013, t(5.91)=3.49), 1 (p=0.0089, t(2.47)=7.62), and 0 neighbors (p=0.043, t(2.04)=4.55). **f,** In learning mice, there are more growing sheaths with 1 neighbor (p=0.0071, t(6.00)=-4.00). **g,** In learning mice, there are more retracting sheaths with 2 (p=0.024, t(4.00)=-3.62) and 1 neighbors (p=0.0050, t(6.00)=-4.31). **h,** Nodes and gaps in myelin that shorten are not modulated by learning. For detailed statistics, see **Supplementary Table 3, Supplementary Figure 3.**

**Supplementary Figure 5.**
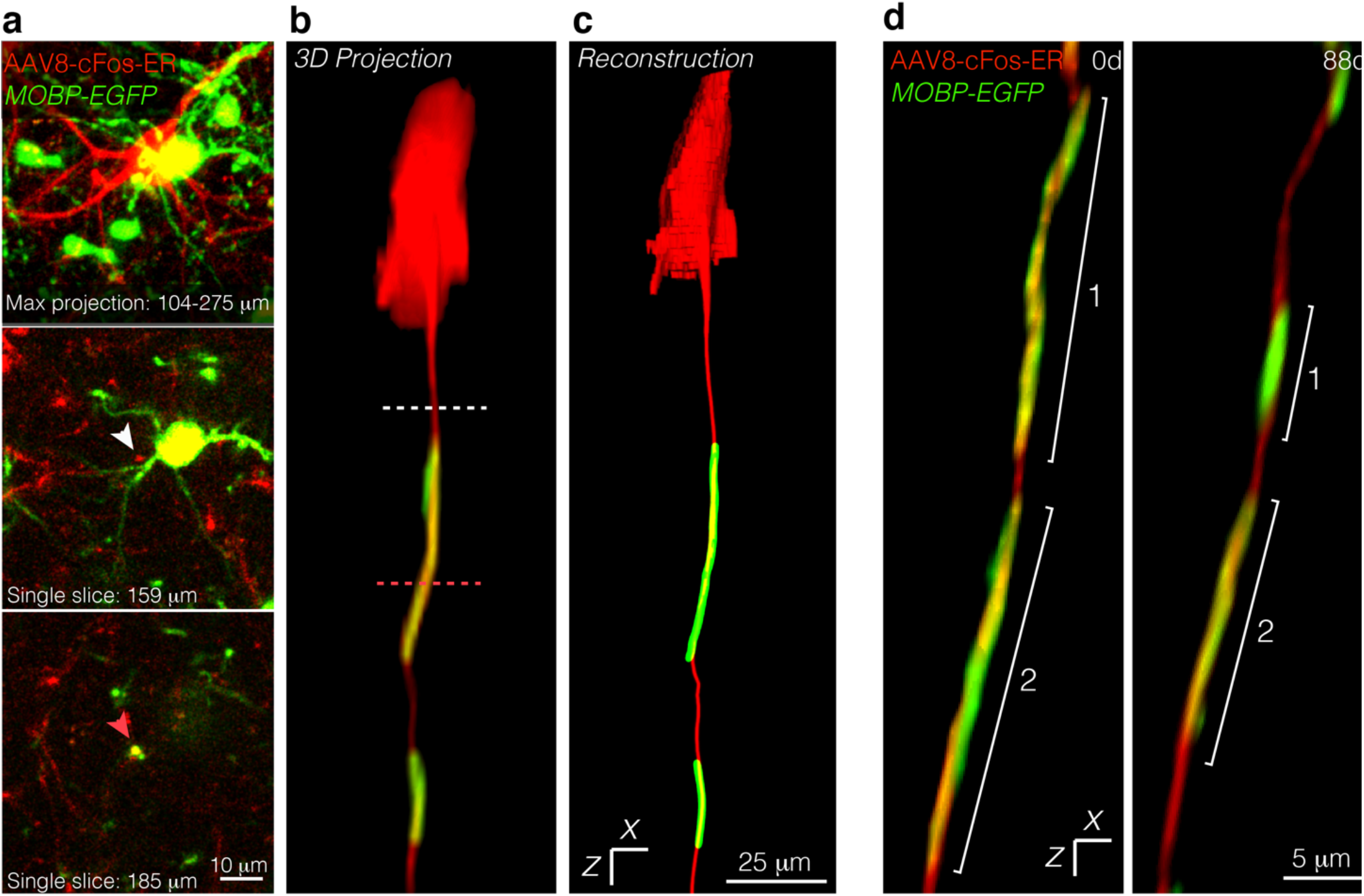
Tracking virally labeled axons across time *in vivo*. **a**, Maximum projection of a cFos+ neuron (AAv8-cFos-ER, red) and surrounding oligodendrocytes and myelin (*in vivo*, green) (top). Single slice at 159 microns below the pial surface, with white arrow identifying unmyelinated region of a cFos+ axon (middle). Single slice at 185 microns below the pial surface, with pink arrow identifying a myelinated region of a cFos+ axon (bottom). **b,** 3D projection of the cFos+ neuron and associated myelin sheaths from (a), with dashed lines corresponding to the single slices in (a). **c,** The same neuron as in (a), reconstructed using Simple Neurite Tracer. **d,** 3D projection of a longitudinally imaged axon at the outset of the imaging experiment (0d) and at the final imaging timepoint (88d).

**Supplementary Figure 6.**
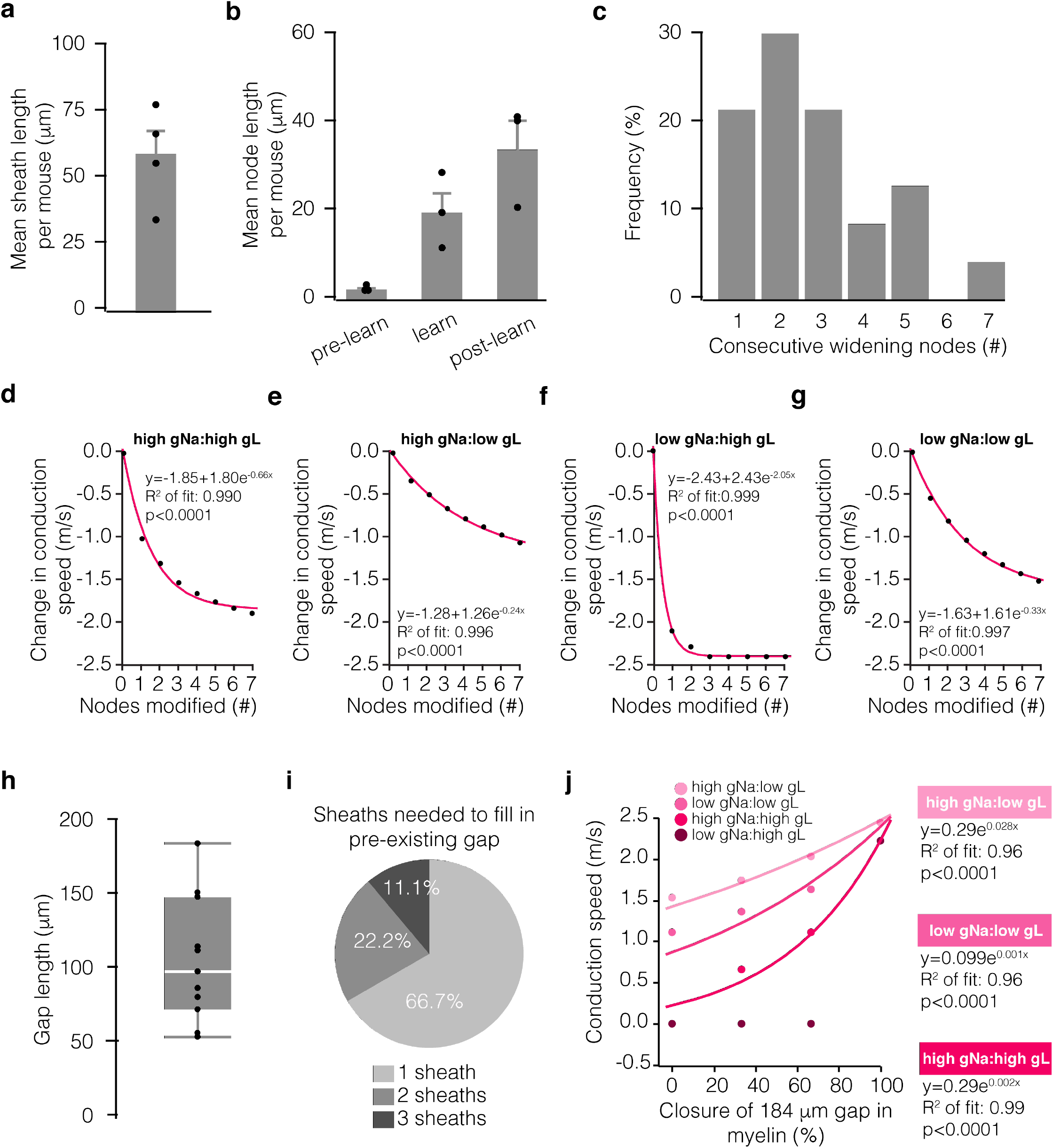
Biological correlates of computational modeling data. **a,** Average pre-existing myelin sheath length per mouse. **b,** Mean node length of lengthening nodes per mouse, measured across time in pre-learning, learning, and post-learning stages. **c,** Distribution of the number of consecutive lengthening nodes along a single axon. **d-g**, Modeled change in conduction speed as a function of the number of consecutively remodeled nodes at different sodium:leak conductance ratios (node with = 35 microns, pink exponential fit). Conductance ratios were generated using either 0.4 S/cm^2^ (low) or 3.4 S/cm^2^ (high) for sodium conductance (gNa) and either 0.01 S/cm^2^ (low) or 0.08S/cm^2^ (high) for leak conductance (gL). **h,** Length of unmyelinated axon in that received new sheath addition following learning, measured from one sheath to the nearest neighboring sheath. Bars and error bars represent median and I.Q.R. **i,** Proportion of gaps filled by 1, 2, or 3 sheaths following learning. **j,** Modeled change in conduction speed as a function of proportion of a 184 micron gap filled in by new myelin (pink exponential fit). Conductance ratios as in **d-g**. Bars and error bars represent mean ± s.e.m unless otherwise noted.

**Supplementary Figure 7.**
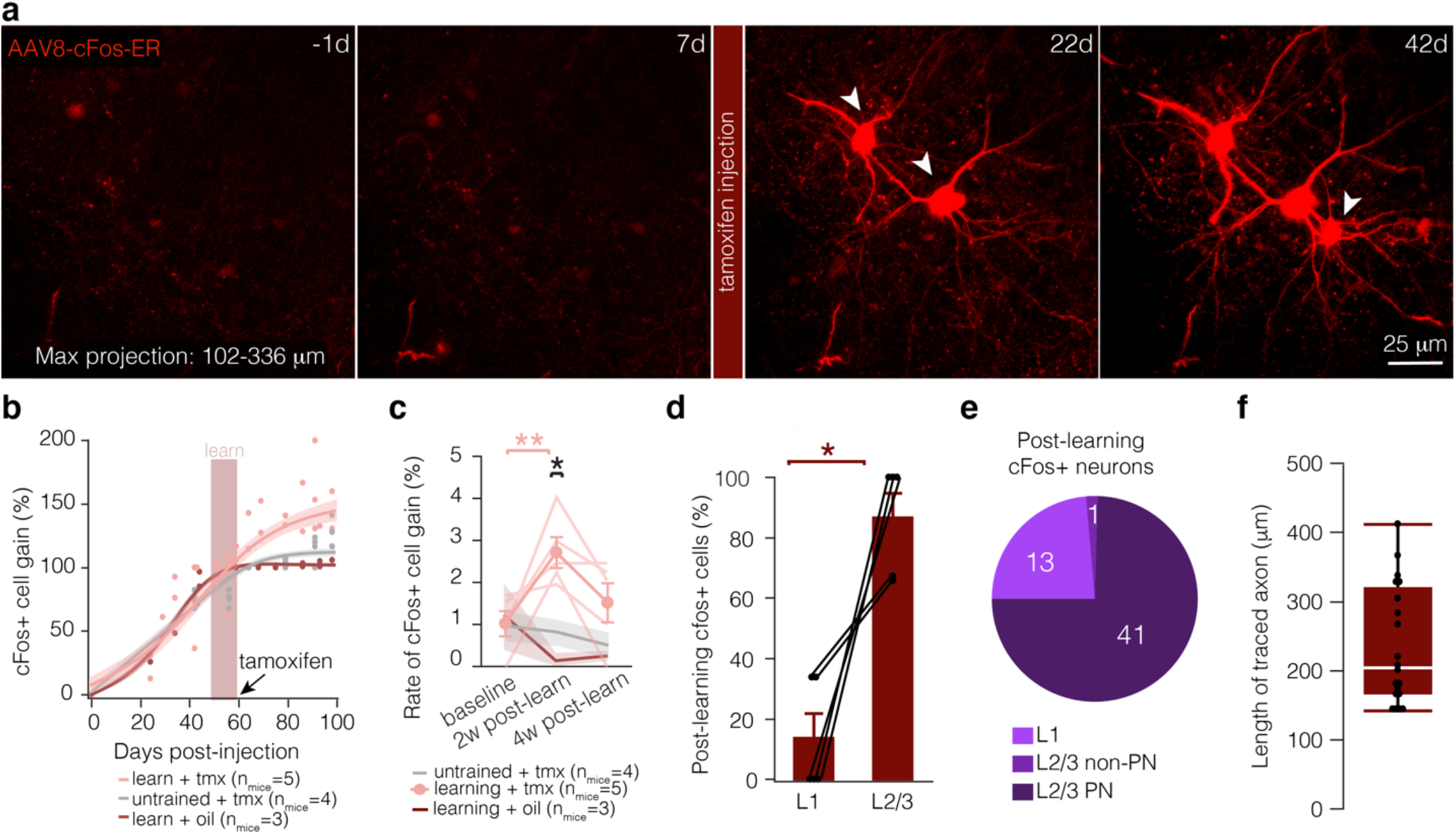
Characterizing AAv8-cFos-ER virus. **a,** Maximum projection of motor cortex injected with the cFos virus 1 day before learning (−1d), one week after learning (7d), three weeks after learning (22d), and 5 weeks after learning (42d). Tamoxifen was injected 3 hours after the final day of learning (7d). **b,** Fit curve for modeled change in cFos+ cell gain (calculated as % of tamoxifen-independent cFos+ neurons, i.e. the number of neurons labeled at 60 days) in mice that learn and receive tamoxifen (pink), untrained mice (grey), and mice that learn and are injected with sunflower oil (red). Each dot represents proportion of labeled neurons per mouse at a given timepoint. **c,** Rate of cFos+ cell gain in mice that learn and receive tamoxifen (pink), untrained mice (grey), and mice that learn and are injected with sunflower oil (red). Mice that learn have a heightened rate of cFos+ cell gain in the two weeks following learning, while the percentage of cFos+ neurons did not change in untrained mice and oil injected mice over the course of the experiment. **d**, Significantly more cFos+ cells appear in L2/3 relative to L1 following learning and injection of tamoxifen. **e,** The majority of learning-activated cells are putative L2/3 pyramidal neurons (determined by morphology). **f,** Distribution of traced axon lengths for cFos+ neurons. For detailed statistics, see **Supplementary Table 3, Supplementary Figure 7.**

