## Supplementary Tables for "Motor Learning Drives Dynamic Patterns of Intermittent Myelination on Learning-activated Axons"

**Supplementary Table 1 | Model Parameters**

| Parameter | Symbol | Value | Units |
|---|---|---|---|
| Nodal membrane capacitance | $C_m$ | 0.9 [1] | $mF/cm^2$ |
| Myelinated axon capacitance | $C_{my}$ | 0.1429 [2] | $mF/cm^2$ |
| Nodal Na+ conductance | $g_{Na}$ | | $S/cm^2$ |
| Lower bound | | 0.0 [3] | |
| Upper bound | | 3.4 [3] | |
| Nodal K+ conductance | $g_K$ | 0.3 | $S/cm^2$ |
| Leak conductance | $g_L$ | | |
| Node | | | $S/cm^2$ |
| Lower bound | | 0. 01 [3] | |
| Upper bound | | 0.08 [1,3] | |
| Myelin sheath | | 0.0001 [1] | $S/cm^2$ |
| Axoplasmic resistivity | $R_a$ | 70 [1] | Ω.cm |
| Resting potential | $E_r$ | -65 | mV |
| Leakage potential | $E_{Lk}$ | -66 | mV |
| Na+ reversal potential | $E_r$ | 55 [2] | mV |
| K+ reversal potential | $E_r$ | -90 [4] | mV |
| Node diameter | | 0.72 [1] | mm |
| Node width | | | mm |
| Lower bound | | 1 * | |
| Upper bound | | 40 * | |
| Myelinated axon diameter | | 0.902 [2,5] | mm |
| g-ratio | | 0.7 [2,5] | |
| Myelin sheath length | | 60 [2, *] | mm |

* Experimentally derived

**Supplementary Table 2 | Antibody specifications**

| | Antigen | Host | Antibody source | Catalog # | Dilution | RRID |
|---|---|---|---|---|---|---|
| Primary antibodies | eGFP | Chicken | AVES | Gfp-1020 | 1:1000 | AB_10000240 |
| | eGFP | Goat | SicGen | AB0020-200 | 1:2000 | AB_2333099 |
| | $Na_v$Pan | Mouse | Neuromab | 75-405 | 1:100 | AB_2491098 |
| | NFH | Chicken | AVES | NFH | 1:1000 | AB_231552 |
| Secondary antibodies | **Antigen** | **Host** | **Antibody source** | **Catalog #** | **Dilution** | **RRID** |
| | Alexa Fluor 488-conjugated anti-chicken IgG | Donkey | Jackson | 703-546-155 | 1:500 | AB_2340376 |
| | Alexa Fluor 488-conjugated anti-goat IgG | Donkey | Jackson | 705-545-003 | 1:500 | AB_2340428 |
| | Alexa Fluor 568-conjugated anti-rabbit IgG | Donkey | Molecular Probes | A10042 | 1:500 | AB_2534017 |
| | Alexa Fluor 568-conjugated anti-guinea pig IgG | Goat | Molecular Probes | A11075 | 1:100 | AB_141954 |
| | Alexa Fluor 647-conjugated anti-chicken IgG | Donkey | Jackson | 103-605-155 | 1:500 | AB_2337392 |

### Supplementary Table 3 | Statistics results table

*N.B. Statistics results are sorted by figure, and both main figures and supplementary figures are listed in the order they appear in the text.*

| FIG1 | Measure | Values | N | Statistical test | Significance |
|---|---|---|---|---|---|
| **Fig 1b** | Cumulative length change (in microns) of individual sheaths with two neighbors (either retracting, stable, or growing) across time (baseline (day -21), pre-learning (day 0), directly post-learning (day 9), two weeks post-learning (day 21) | | **Growing sheaths:** n=48 sheaths, n=4 mice<br><br>**Stable sheaths:** n=24 sheaths, n=4 mice<br><br>**Retracting sheaths:** n=52 sheaths, n=4 mice | ANOVA<br>**Growing sheaths:** F(3,44)=7.24, p=0.0005<br><br>**Stable sheaths:** F(3,20)=0.2194, p=0.88<br><br>**Retracting sheaths:** F(3,48)=9.04, p<0.0001<br><br>Post-hoc tests using Tukey's HSD | **Growing sheaths**:<br>Day -21 vs. day 0: p=0.001<br><br>Day -21 vs. day 9: p=0.0015<br><br>Day -21 vs. day 21: p=0.0211<br><br>**Retracting sheaths**:<br>Day -21 vs. day 21: p=0.0016<br><br>Day 0 vs. day 21: p<0.0001<br><br>Day 9 vs. day 21: p=0.0213<br><br>All other comparisons p>0.05 |
| **Fig 1c** | Cumulative length change (in microns) of individual sheaths with one neighbor (either retracting, stable, or growing) across time (baseline (day -21), pre-learning (day 0), directly post-learning (day 9), two weeks post-learning (day 21) | | **Growing sheaths:** n=59 sheaths, n=4 mice<br><br>**Stable sheaths:** n=24 sheaths, n=4 mice<br><br>**Retracting sheaths:** n=43 sheaths, n=4 mice | ANOVA<br>**Growing sheaths:** F(3,56)=4.46, p=0.007<br><br>**Stable sheaths:** F(3,20)=0.22, p=0.88<br><br>**Retracting sheaths:** F(3,40)=9.93, p<0.0001<br><br>Post-hoc tests using Tukey's HSD | **Growing sheaths**:<br>Day -21 vs. day 21: p=0.0043<br><br>**Retracting sheaths**:<br>Day -21 vs. day 21: p=0.0002<br><br>Day -21 vs. day 9: p=0.0012<br><br>Day 0 vs. day 9: p=0.029<br><br>Day 0 vs. day 21: p=0.0071<br><br>All other comparisons p>0.05 |
| **Fig. 1d** | Cumulative length change (in microns) of individual sheaths with 0 neighbors (either retracting, stable, or growing) across time (baseline (day -21), pre-learning (day 0), directly post-learning (day 9), two weeks post-learning (day 21) | | **Growing sheaths:** n=27 sheaths, n=4 mice<br><br>**Stable sheaths:** n=19 sheaths, n=4 mice<br><br>**Retracting sheaths:** n=43 sheaths, n=4 mice | ANOVA<br>**Growing sheaths:** F(3,24)=3.97, p=0.020<br><br>**Stable sheaths:** F(3,19)=0.36, p=0.79<br><br>**Retracting sheaths:** F(3,40)=3.08, p=0.038<br><br>Post-hoc tests using Tukey's HSD | **Growing sheaths**:<br>Day -21 vs. day 0: p=0.028<br><br>Day -21 vs. day 21: p=0.032<br><br>**Retracting sheaths**:<br>Day -21 vs. day 21: p=0.025<br><br>All other comparisons p>0.05 |

| | | | | | |
|---|---|---|---|---|---|
| **Fig. 1f** | Proportion of nodes engaging in widening (% of all nodes) pre-learning and post-learning in learning mice compared to age-matched controls (mean±SEM) | **Pre-learn:**<br>Untrained: 3.44±1.36<br>Learning: : 1.82±80<br><br>**Post-learn:**<br>Untrained: 3.06±1.00<br>Learning: 21.73±2.60 | **Untrained:**<br>n=312 nodes, n=6 mice<br><br>**Learning:**<br>n=227 nodes, n=5 mice | Restricted Maximum Likelihood model (REML) to predict node width with full factorial model between training (untrained vs learning) and day (pre-learning vs. post-learning), and with random variable of Mouse.<br><br>Interaction effect between Training and Day to predict node width: F(1,14)=438.85, p=0.0002<br><br>$R^2$=0.79 | All post-hoc tests conducted using Tukey's HSD:<br><br>**Learning post-learn** vs. **Untrained pre-learn**: p<0.0001<br><br>**Learning post-learn** vs. **Untrained post-learn**: p<0.0001<br><br>**Learning pre-learn** vs. **Learning post-learn:** p<0.0001<br><br>All other relevant comparisons p>0.05 |
| **Fig. 1g** | Node width (in microns) of individual widening nodes across time (baseline (d -21), pre-learning (d1), directly post-learning (d 9), two weeks post-learning (d 21) in learning mice compared to age-matched controls (mean±SEM) | **Baseline:**<br>Untrained: 1.67±0.16<br>Learning: : 2.07±0.19<br><br>**Pre-learning:**<br>Untrained: 2.41±0.96<br>Learning: 5.37±2.03<br><br>**Directly post-learning:**<br>Untrained: 3.63±0.92<br>Learning: 18.72±4.31<br><br>**Two weeks post-learning:**<br>Untrained: 6.02±1.28<br>Learning: 32.43±5.25 | **Untrained:**<br>n=27 nodes, n=3 mice<br><br>**Learning:**<br>n=67 nodes, n=4 mice | Restricted Maximum Likelihood model (REML) to predict node width with full factorial model between training (untrained vs learning) and day (baseline vs. pre-learning vs. directly post-learn vs. two weeks post-learning) and with random variable of Mouse.<br><br>Interaction effect between Training and Day to predict node width: F(3,83.38)=4.88, p=0.0036 | All post-hoc tests conducted using Tukey's HSD:<br><br>**Learning d-21** vs. **Learning d9**: p=0.0014<br>**Learning d-21** vs. **Learning d21:**p<0.0001<br>**Learning d1** vs. **Learning d9**: p=0.039<br>**Learning d1** vs. **Learning d21:**p<0.0001<br>**Learning d9** vs. **Learning d21:** p=0.030<br><br>**Untrained d21 vs. Learning d21**: p=0.015<br><br>All other relevant comparisons p>0.05 |
| **Fig. 1h** | Relationship between the cumulative sheath retraction and change in associated node width | | **n=32 paranodes/16 associated nodes** | Standard least squares to predict change in node width using cumulative sheath retraction<br><br>R-square = 0.819, p<0.0001 | |

| **SUPP. FIG2** | **Measure** | **Values** | **N** | **Statistical test** | **Significance** |
|---|---|---|---|---|---|
| **Supp. Fig. 2d** | Change in success rate on the first and last day of learning | **Learning**<br>First: 12.25±1.08<br>Last: 44.17±5.04 | **Learning**<br>n=12 mice | Paired student's t-test<br><br>**Learning**<br>t(11)=-2.78 | **Learning**<br>p<0.0001<br>Cohen's d = 1.66<br><br>All other relevant comparisons p>0.05 |
| **Supp. Fig. 2e** | Success rate (%) across all figures at Day 1 of learning and Day 7 of learning | **Fig 1**<br>Day 1: 10.5 ± 4.59<br>Day 7: 46.6 ± 6.66<br>**Fig 2**<br>Day 1: 12.6 ± 3.81<br>Day 7: 52.7 ± 3.39<br>**Fig 3**<br>Day 1: 10.5 ± 3.75<br>Day 7: 47.8 ± 5.58 | **Fig 1**<br>n = 5 mice<br>**Fig 2**<br>n = 5 mice<br>**Fig 3**<br>n = 6 mice<br>**Fig 4**<br>n = 9 mice<br>**Fig 6** | Restricted Maximum Likelihood model (REML) to predict success rate (%) versus training day (Day 1 vs. Day 7) and with random variable of Mouse.<br><br>F(6,6)=22.57, p=1.0 | All relevant comparisons p>0.05 |

| | | | | | |
|---|---|---|---|---|---|
| | | **Fig 4**<br>Day 1: 9.42 ± 2.79<br>Day 7: 46.8 ± 4.27<br>**Fig 6**<br>Day 1: 16.8 ± 1.79<br>Day 7: 56.4 ± 2.03<br>**Fig 7**<br>Day 1: 6.4 ± 2.1<br>Day 7: 32.6 ± 4.64 | n = 5 mice<br>**Fig 7**<br>n = 14 mice | | |
| **Supp. Fig. 2f** | Change in success rate (%) across all figures at Day 1 of learning and Day 7 of learning | **Fig 1**<br>36.1 ± 4.28<br>**Fig 2**<br>40.1 ± 3.06<br>**Fig 3**<br>37.4 ± 3.72<br>**Fig 4**<br>37.4 ± 2.58<br>**Fig 6**<br>39.6 ± 2.96<br>**Fig 7**<br>26.2 ± 3.51 | **Fig 1**<br>n = 5 mice<br>**Fig 2**<br>n = 5 mice<br>**Fig 3**<br>n = 6 mice<br>**Fig 4**<br>n = 9 mice<br>**Fig 6**<br>n = 5 mice<br>**Fig 7**<br>n = 14 mice | Restricted Maximum Likelihood model (REML) to predict change in success rate (%) with random variable of Mouse.<br><br>F(5,5)=13.53, p=1.0 | All relevant comparisons p>0.05 |
| **SUPP. Fig3** | **Measure** | **Values** | **N** | **Statistical test** | **Significance** |
| **Supp. Fig. 3c** | Proportion of nodes in fixed tissue vs. nodes identified in longitudinal images with NaV (%) | **Fixed tissue**<br>98.05 ± 1.94<br><br>**Live imaged**<br>90.9 ± 6.46 | **Fixed tissue**<br>n=6 mice<br><br>**Live imaged**<br>n=3 mice | Student's t-test<br><br>t(-1.87)=2.13 | p>0.05<br><br>Cohen's d = 1.5 |
| | Proportion of gaps with and without NaV (%) | **NaV present**<br>52.3 ± 4.54<br><br>**NaV absent**<br>47.7 ± 4.54 | n=8 mice | Paired student's t-test<br><br>t(-1.87)=2.13 | p>0.05<br><br>Cohen's d = 1.5 |
| **SUPP. Fig4** | **Measure** | **Values** | **N** | **Statistical test** | **Significance** |
| **Supp Fig. 4a** | Sheath length (um) in live-imaged sheaths with 2 vs. 1 vs. 0 neighbors | **2 neighbors**<br>59.5 ± 9.33<br><br>**1 neighbor**<br>52.9 ± 7.21<br><br>**0 neighbors**<br>59.5 ± 7.94 | n = 4 (2 neighbors) or 3 (1 or 0 neighbors) mice | ANOVA<br>**Growing sheaths:**<br>F(2,7)=0.1812, p > 0.8 | |
| **Supp Fig. 4c** | Proportion (%) of all dynamic sheaths growing pre-, during, and post-learning in untrained vs. learning mice | **Pre-learn:**<br>Learning: 46.3 ± 10.9<br>Untrained: 39.6 ± 6.25<br><br>**Learn:**<br>Learning: 19.2 ± 7.98<br>Untrained: 25.0 ± 3.4<br><br>**Post-learn:**<br>Learning: 27.3 ± 10.6<br>Untrained: 35.4 ± 5.24 | n=4 mice<br>n=5 mice | Restricted Maximum Likelihood model (REML) to predict sheath growth with full factorial model between training (untrained vs learning) and stage (pre-learn vs. learn vs. post-learn) and with random variable of Mouse.<br><br>Interaction effect between Training and Day to predict sheath growth: F(3, 9.46)=16.14, p>0.05<br><br>$R^2$=0.64 | All post-hoc tests conducted using Tukey's HSD:<br><br>All comparisons p>0.05 |
| **Supp Fig. 4d** | Proportion (%) of all dynamic sheaths retracting pre-, during, and post-learning in | **Pre-learn:**<br>Learning: 29.6 ± 4.78<br>Untrained: 34.4 ± 6.88 | n=4 mice<br>n=5 mice | Restricted Maximum Likelihood model (REML) to predict sheath retraction with full factorial model between | All post-hoc tests conducted using Tukey's HSD: |

| | | | | | |
|---|---|---|---|---|---|
| | untrained vs. learning mice | **Learn:**<br>Learning: 64.3 ± 7.64<br>Untrained: 22.9 ± 10.4<br><br>**Post-learn:**<br>Learning: 60.3 ± 6.34<br>Untrained: 28.1 ± 4.62 | | training (untrained vs learning) and stage (pre-learn vs. learn vs. post-learn) and with random variable of Mouse.<br><br>Interaction effect between Training and Day to predict sheath retraction: F(3, 115.8)=6.76, p=0.0088<br><br>$R^2$=0.83 | **Untrained, learn,** vs. **Learning, learn**: p=0.0095<br><br>**Learning, pre-learn** vs. **Learning, learn**: p=0.016<br><br>**Learning, pre-learn** vs. **Learning, post-learn**: p=0.0358<br><br>All other relevant comparisons p>0.05 |
| **Supp Fig. 4e** | Proportion of sheaths with 2, 1, and 0 neighbors that remain stable across the experiment in learning vs. untrained mice (%) | **2 neighbors:**<br>Learning: 86.5 ± 2.08<br>Untrained: 98.2 ± 1.75<br>**1 neighbor:**<br>Learning: 37.8 ± 4.74<br>Untrained: 82.6 ± 3.24<br>**0 neighbors:**<br>Learning: 44.6 ± 6.72<br>Untrained: 77.6 ± 1.87 | **Learning:**<br>n = 5 mice<br><br>**Untrained:**<br>n=3 mice | Student's t-test<br><br>**2 neighbors:**<br>t(3.49)=5.91<br><br>**1 neighbor:**<br>t(2.47)=7.61<br><br>**0 neighbors:**<br>t(4.55)=2.04 | **2 neighbors:**<br>p=0.13<br>Cohen's d = 6.09<br><br>**1 neighbor:**<br>p=0.0089<br>Cohen's d = 11.03<br><br>**0 neighbors:**<br>p=0.44<br>Cohen's d = 6.69 |
| **Supp Fig. 4f** | Proportion of sheaths with 2, 1, and 0 neighbors that grow across the experiment in learning vs. untrained mice (%) | **2 neighbors:**<br>Learning: 3.25 ± 2.08<br>Untrained: 1.75 ± 1.75<br>**1 neighbor:**<br>Learning: 35.9 ± 4.74<br>Untrained: 12.9 ± 3.24<br>**0 neighbors:**<br>Learning: 18.3 ± 6.72<br>Untrained: 3.72 ± 1.87 | **Learning:**<br>n = 5 mice<br><br>**Untrained:**<br>n=3 mice | Student's t-test<br><br>**2 neighbors:**<br>t(5.82)=-0.55<br><br>**1 neighbor:**<br>t(6.00)=-4.00<br><br>**0 neighbors:**<br>t(4.58)=-2.09 | **2 neighbors:**<br>p>0.6<br><br>**1 neighbor:**<br>p=0.0071<br>Cohen's d = 5.67<br><br>**0 neighbors:**<br>p>009 |
| **Supp Fig. 4g** | Proportion of sheaths with 2, 1, and 0 neighbors that retract across the experiment in learning vs. untrained mice (%) | **2 neighbors:**<br>Learning: 10.2 ± 2.82<br>Untrained: 0.00 ± 0.00<br>**1 neighbor:**<br>Learning: 25.5 ± 4.01<br>Untrained: 4.51 ± 2.76<br>**0 neighbors:**<br>Learning: 37.1 ± 6.36<br>Untrained: 18.7 ± 6.73 | **Learning:**<br>n = 5 mice<br><br>**Untrained:**<br>n=3 mice | Student's t-test<br><br>**2 neighbors:**<br>t(4)=-3.62<br><br>**1 neighbor:**<br>t t(6.00)=-4.31<br><br>**0 neighbors:**<br>t(5.12)=-1.98 | **2 neighbors:**<br>p=0.02<br>Cohen's d = 5.11<br><br>**1 neighbor:**<br>p=0.0050<br>Cohen's d = 6.10<br><br>**0 neighbors:**<br>p>0.10 |
| **Supp Fig. 4h** | Change in node width (in microns) of individual shrinking nodes across time (baseline (d -21), pre-learning (d1), directly post-learning (d 9), two weeks post-learning (d 21) in learning mice compared to age-matched controls (mean±SEM) | **Baseline:**<br>Untrained: 0.00±0.00<br>Learning: : 0.00±0.00<br><br>**Pre-learning:**<br>Untrained: -3.34 ± 2.06<br>Learning: -2.72 ± 1.69<br><br>**Directly post-learning:**<br>Untrained: -3.21 ± 1.83<br>Learning: -6.79 ± 2.79 | **Untrained:**<br>n=15 nodes, n=3 mice<br><br>**Learning:**<br>n=30 nodes, n=4 mice | Restricted Maximum Likelihood model (REML) to predict node width with full factorial model between training (untrained vs learning) and day (baseline vs. pre-learning vs. directly post-learn vs. two weeks post-learning) and with random variable of Mouse.<br><br>Interaction effect between Training and Day to predict node width: F(1,37.05)=-.13, p>0.07 | |

| | | | | | |
|---|---|---|---|---|---|
| | | **Two weeks post-learning:**<br>Untrained: -6.52 ± 1.24<br>Learning: -7.84 ± 3.57 | | | |
| **FIG2** | **Measure** | **Values** | **N** | **Statistical test** | **Significance** |
| **Fig 2b** | Number of sheaths per new oligodendrocyte in learning mice and age-matched controls (mean±SEM) | **Untrained:** 38.25±3.30<br><br>**Learning:** 44.33±4.04 | n=4 untrained mice<br>n = 3 learning mice | Student's t-test<br><br>t(3.86)=-2.12 | p>0.10 |
| **Fig 2c** | Sheath length (in microns) of individual new myelin sheaths in learning mice and age-matched controls<br><br>Values are shown as median (IQR). | **Untrained:** 57.58 (15.07) microns<br><br>**Learning:** 47.72 (24.43) microns | Untrained, n= 20 sheaths<br><br>Learning, n=49 sheaths | Student's t-test<br><br>t(244.18)=1.19 | p>0.24 |
| **Fig 2d** | Proportion of new sheaths with 2, 1, and 0 neighbors per new oligodendrocyte (%) in learning mice and age-matched controls (mean±SEM) | **Untrained**<br>2 neighbors: 9.15±2.62<br>1 neighbor: 30.00±4.75<br>0 neighbors: 60.85±6.97<br><br>**Learning:**<br>2 neighbors: 30.66±3.54<br>1 neighbor: 30.32±4.01<br>0 neighbors: 39.02±0.53 | n=4 untrained mice<br>n = 3 learning mice | Two-way ANOVA between number of neighbors (2 vs. 1 vs. 0) and training (untrained versus learning)<br><br>F(5,15)=15.03, p<0.0001<br><br>All post-hoc tests using Tukey's HSD | Within group effects:<br>**Untrained,**<br>2 vs 1 neighbor: p=0.036<br>2 vs 0 neighbors: p<0.0001<br>1 vs 0 neighbors: p=0.0016<br><br>Between-group effects:<br>**2 neighbors,** untrained vs. learning: p=0.048<br>**0 neighbors,** untrained vs. learning: p=0.044<br><br>All other relevant comparisons p>0.05 |
| **Fig 2f** | Proportion of new sheaths (%) filling in gaps in pre-existing myelin in learning mice compared to age-matched controls (mean±SEM) | **Untrained:** 6.70±3.88%<br><br>**Learning:** 32.43±8.38% | n=4 untrained mice<br><br>n=6 learning mice | Student's t-test<br><br>t(6.85)=-2.78 | p=0.028<br>Cohen's d = 1.66 |
| **Fig 2g** | Proportion of gaps in myelin filled in by new sheaths (%) that are pre-existing gaps in myelin vs. widened nodes in learning mice (mean±SEM) | **Widened:** 10.67±6.86%<br><br>**Pre-existing:** 91.11±5.88% | n=6 learning mice | Paired student's t-test<br><br>t(5)=6.99 | p=0.0009<br>Cohen's d=5.71 |
| **Fig 2i** | Proportion of axons (%) with node and myelin dynamics in learning vs. untrained mice | **Untrained:** 14.2 ± 4.79%<br><br>**Learning:** 60.3 ± 11.4% | n=4 untrained mice<br><br>n=6 learning mice | Student's t-test:<br><br>t(6.57)=-3.73 | p=0.0082<br>Cohen's d=5.27 |
| **Fig 2k** | Timing (in days) of pre-existing sheath retraction versus new sheath addition | **Sheath retraction:** 6.47±1.87 days | **Sheath retraction:** n=8 learning mice | Student's t-test:<br><br>t(13.43)=-2.46 | p=0.028<br>Cohen's d=1.23 |

| | | | | | |
|---|---|---|---|---|---|
| | relative to the onset of learning (mean±SEM) | **Sheath addition:** 12.39±1.51 days | **Sheath addition:** n=8 learning mice | | |
| **FIG. 3** | **Measure** | **Values** | **N** | **Statistical test** | **Significance** |
| **Fig 3c** | Proportion of axons with dynamic myelin (%) that show majority stable vs. majority dynamic sheaths along their length in learning mice vs. age-matched controls (mean±SEM) | **Untrained**<br>Majority stable: 90.83±9.41<br>Majority dynamic: 9.16±9.41<br><br>**Learning:**<br>Majority stable: 31.11±12.14<br>Majority dynamic: 68.89±12.14 | n=5 untrained mice<br>n = 3 learning mice | Two-way ANOVA between axon-specific sheath behavior (majority stable vs. majority dynamic) and training (untrained vs. learning)<br><br>$F(3,12)=14.17$, $p=0.0003$<br><br>All post-hoc tests using Tukey's HSD | Within group effects:<br>**Untrained,** majority stable vs. majority dynamic: $p=0.0003$<br><br>Between-group effects:<br>**Majority stable,** untrained vs. learning: $p=0.01$<br>**Majority dynamic,** untrained vs. learning: $p=0.01$<br><br>All other relevant comparisons $p>0.05$ |
| **Fig 3e** | Proportion of axons (%) showing heightened sheath addition pre- and post-learning in learning mice compared to age-matched controls (mean±SEM) | **Pre-learn:**<br>Untrained: 19.95±3.31<br>Learning: : 19.83±3.63<br><br>**Post-learn:**<br>Untrained: 16.79±3.62<br>Learning: 41.23±4.06 | **Untrained:** n=6 mice<br><br>**Learning:** n=5 mice | Restricted Maximum Likelihood model (REML) to predict heightened sheath addition with full factorial model between training (untrained vs learning) and stage (pre-learn vs. post-learn) and with random variable of Mouse.<br><br>Interaction effect between Training and Day to predict node width: $F(1, 9.46)=16.14$, $p=0.0027$<br><br>$R^2=0.81$ | All post-hoc tests conducted using Tukey's HSD:<br><br>**Untrained, post-learn** vs. **Learning, post-learn**: $p=0.006$<br><br>**Untrained, pre-learn** vs. **Learning, post-learn**: $p=0.011$<br><br>**Learning, pre-learn** vs. **Learning, post-learn**: $p=0.0055$<br><br>All other relevant comparisons $p>0.05$ |
| **FIG 4** | **Measure** | **Values** | **N** | **Statistical test** | **Significance** |
| **Fig. 4b** | Proportion of Layer 1 nodes engaging in widening (% of all nodes) pre-learning and post-learning in learning mice compared to age-matched controls (mean±SEM) | **Pre-learn:**<br>Untrained: 1.88±0.77<br>Learning: : 1.11±0.84<br><br>**Post-learn:**<br>Untrained: 1.36±0.59<br>Learning: 19.84±2.09 | **Untrained:** n=286 nodes, n=8 mice<br><br>**Learning:** n=286 nodes, n=8 mice | Restricted Maximum Likelihood model (REML) to predict node width with full factorial model between training (untrained vs learning) and day (pre-learning vs. post-learning) and with random variable of Mouse.<br><br>Interaction effect between Training and Day to predict node width: $F(1,14)=49.33$, $p<0.0001$<br><br>$R^2=0.74$ | All post-hoc tests conducted using Tukey's HSD:<br><br>**Learning post-learn** vs. **Untrained pre-learn**: $p<0.0001$<br><br>**Learning post-learn** vs. **Untrained post-learn**: $p<0.0001$<br><br>**Learning pre-learn** vs. **Learning post-learn:** $p<0.0001$<br><br>All other relevant comparisons $p>0.05$ |
| **Fig. 4c** | Proportion of L1 axons with dynamic myelin (%) that show majority stable vs. majority dynamic sheaths along their | **Untrained**<br>Majority stable: 90.28±6.24<br>Majority dynamic: 9.72±6.24 | n=6 untrained mice<br>n = 8 learning mice | Two-way ANOVA between axon-specific sheath behavior (majority stable vs. majority dynamic) and training (untrained vs. learning) | Within group effects:<br>**Untrained,** majority stable vs. majority dynamic: $p<0.0001$<br>**Learning,** |

| | | | | | |
|---|---|---|---|---|---|
| | length in learning mice vs. age-matched controls (mean±SEM) | **Learning:**<br>Majority stable: 33.33±6.86<br>Majority dynamic: 66.67±6.86 | | $F(3,24)=25.12$, $p<0.0001$<br><br>$R^2=0.73$<br><br>All post-hoc tests using Tukey's HSD | majority stable vs. majority dynamic: p=0.0052<br><br>Between-group effects:<br>**Majority stable,** untrained vs. learning: p<0.0001<br>**Majority dynamic,** untrained vs. learning: p<0.0001<br><br>All other relevant comparisons p>0.05 |
| **Fig. 4d** | Proportion of L1 axons (%) showing heightened sheath addition pre- and post-learning in learning mice compared to age-matched controls (mean±SEM) | **Pre-learn:**<br>Untrained: 21.07±2.83<br>Learning: : 21.59±2.01<br><br>**Post-learn:**<br>Untrained: 19.64±1.72<br>Learning: 39.46±5.97 | **Untrained:** n=9 mice<br><br>**Learning:** n=7 mice | Restricted Maximum Likelihood model (REML) to predict heightened sheath addition with full factorial model between training (untrained vs learning) and stage (pre-learn vs. post-learn) and with random variable of Mouse.<br><br>Interaction effect between Training and Day to predict node width: $F(1,11.95)=12.65$, $p=0.0040$<br><br>$R^2=0.82$ | All post-hoc tests conducted using Tukey's HSD:<br><br>**Untrained, post-learn** vs. **Learning, post-learn**: p=0.0087<br><br>**Untrained, pre-learn** vs. **Learning, post-learn**: p=0.015<br><br>**Learning, pre-learn** vs. **Learning, post-learn**: p=0.0065<br><br>All other relevant comparisons p>0.05 |
| **Fig. 4e** | Timing (in days) of pre-existing sheath retraction versus new sheath addition relative to the onset of learning in Layer 1 (mean±SEM) | **Sheath retraction:** 7.02±0.88 days<br><br>**Sheath addition:** 11.29±0.36 days | **Sheath retraction:** n=4 learning mice<br><br>**Sheath addition:** n=4 learning mice | Student's t-test: $t(4.00)=-4.51$ | p=0.011<br>Cohen's d=3.19 |
| **Fig. 4h** | Proportion of Layer 2/3 nodes engaging in widening (% of all nodes) pre-learning and post-learning in learning mice compared to age-matched controls (mean±SEM) | **Pre-learn:**<br>Untrained: 4.30±2.41<br>Learning: : 2.82±1.84<br><br>**Post-learn:**<br>Untrained: 5.48±2.56<br>Learning: 26.11±2.53 | **Untrained:** n=107 nodes, n=6 mice<br><br>**Learning:** n=65 nodes, n=5 mice | Restricted Maximum Likelihood model (REML) to predict node width with full factorial model between training (untrained vs learning) and day (pre-learning vs. post-learning) and with random variable of Mouse.<br><br>Interaction effect between Training and Day to predict node width: $F(1,9)=15.85$, $p=0.0032$<br><br>$R^2=0.45$ | All post-hoc tests conducted using Tukey's HSD:<br><br>**Learning post-learn** vs. **Untrained pre-learn**: p=0.0006<br><br>**Learning post-learn** vs. **Untrained post-learn**: p=0.0009<br><br>**Learning pre-learn** vs. **Learning post-learn:** p=0.0014<br><br>All other relevant comparisons p>0.05 |
| **Fig. 4i** | Proportion of L2/3 axons with dynamic myelin (%) that show majority stable vs. majority dynamic sheaths along their length in learning | **Untrained**<br>Majority stable: 85.42±8.59<br>Majority dynamic: 14.58±8.59<br><br>**Learning:** | n=4 untrained mice<br>n = 4 learning mice | Two-way ANOVA between axon-specific sheath behavior (majority stable vs. majority dynamic) and training (untrained vs. learning)<br><br>$F(3,12)=9.87$, $p=0.0015$ | Within group effects:<br>**Untrained,** majority stable vs. majority dynamic: p=0.0021<br><br>Between-group effects: |

| | | | | | |
|---|---|---|---|---|---|
| | mice vs. age-matched controls<br>(mean±SEM) | Majority stable:<br>31.25±11.97<br>Majority dynamic:<br>68.75±11.97 | | $R^2$=0.64<br><br>All post-hoc tests using Tukey's HSD | **Majority stable,** untrained vs. learning:<br>p=0.0145<br>**Majority dynamic,** untrained vs. learning:<br>p=0.0145<br><br>All other relevant comparisons p>0.05 |
| **Fig. 4j** | Proportion of L2/3 axons (%) showing heightened sheath addition pre- and post-learning in learning mice compared to age-matched controls (mean±SEM) | **Pre-learn:**<br>Untrained: 6.89±2.86<br>Learning: : 5.79±3.87<br><br>**Post-learn:**<br>Untrained: 7.50±4.79<br>Learning: 32.27±7.11 | **Untrained:**<br>n=5 mice<br><br>**Learning:** n=6 mice | Restricted Maximum Likelihood model (REML) to predict heightened sheath addition with full factorial model between training (untrained vs learning) and stage (pre-learn vs. post-learn) and with random variable of Mouse.<br><br>Interaction effect between Training and Day to predict node width: F(1,8.92)=7.87, p=0.021<br><br>$R^2$=0.70 | All post-hoc tests conducted using Tukey's HSD:<br><br>**Untrained, post-learn** vs. **Learning, post-learn**: p=0.048<br><br>**Untrained, pre-learn** vs. **Learning, post-learn**: p=0.027<br><br>**Learning, pre-learn** vs. **Learning, post-learn**: p=0.0063<br><br>All other relevant comparisons p>0.05 |
| **Fig. 4k** | Timing (in days) of pre-existing sheath retraction versus new sheath addition relative to the onset of learning in Layer 2/3 (mean±SEM) | **Sheath retraction:**<br>4.92±1.33 days<br><br>**Sheath addition:**<br>12.50±2.25 days | **Sheath retraction:** n=4 learning mice<br><br>**Sheath addition:** n=4 learning mice | Student's t-test: t(4.88)=-2.91 | p=0.035<br>Cohen's d=2.05 |
| **FIG. 5** | **Measure** | **Values** | **N** | **Statistical test** | **Significance** |
| **Fig. 5g** | Comparison of change in conduction speed relative to 2.44 m/s (conduction speed on a control axon) as consecutive nodes are widened to 35 microns in concert (0 vs. 1 vs. 2 vs. 3 vs. 4 vs. 5 vs. 6 vs. 7 nodes widened at once) (mean±SEM) | **0 nodes modified:**<br>0.00±0.00<br><br>**1 node modified:**<br>-1.00±0.40<br><br>**2 nodes modified:**<br>-1.23±0.40<br><br>**3 nodes modified:**<br>-1.41±0.39<br><br>**4 nodes modified:**<br>-1.51±0.36<br><br>**5 nodes modified:**<br>-1.59±0.33<br><br>**6 nodes modified:**<br>-1.66±0.31<br><br>**7 nodes modified**<br>-1.72±0.29 | **Number of conditions (defined as different gNa:gL ratios):** n=4 conditions | Restricted Maximum Likelihood model (REML) to predict conduction speed (m/s) with random variable of gNa:gL ratio<br><br>F(7,21)=17.33, p<0.0001 | **Node 0 vs. Node 1**<br>p=0.0008<br><br>**Node 0 vs. Node 2,3,4,5,6,7**<br>p<0.0001<br><br>**Node 1 vs. Node 6**:<br>p=0.042<br><br>**Node 1 vs. Node 7**:<br>p=0.021<br><br>All other comparisons p>0.05 |
| **Fig. 5j** | Conduction speed (m/s) as a function of number of new sheaths added (#) in four conditions, ie.e gNa:gL ratios (high:low, low:low, | **0 sheaths added:**<br>0.66±0.39<br><br>**1 sheath added:**<br>0.94±0.39<br><br>**2 sheaths added:** | **Number of conditions (defined as different gNa:gL ratios):** n=4 conditions | Restricted Maximum Likelihood model (REML) to predict conduction speed (m/s) with random variable of gNa:gL ratio<br><br>F(3,9)=14.39, p=0.0009 | **3 sheaths vs. 0 sheaths added**:<br>p=0.0008<br><br>**3 sheaths vs. 1 sheath added**:<br>p=0.0029 |

| | | | | | |
|---|---|---|---|---|---|
| | low:high, and high:high) | 1.19±0.44<br><br>**3 sheaths added:** 2.39±0.06 | | | **3 sheaths vs. 2sheaths added**: p=0.01<br><br>All other comparisons p>0.05 |
| **FIG 6** | **Measure** | **Values** | **N** | **Statistical test** | **Significance** |
| **Fig. 6d** | Number of dynamic pre-existing sheaths per axon (#) on learning-activated axons compared to axons in age-matched untrained controls and baseline-labeled axons in the same mice. (mean±SEM) | **Untrained axons:** 0.00 ± 0.00<br><br>**Baseline-labeled axons:** 0.533 ± 0.37<br><br>**Learning-activated axons:** 2.3 ± 0.30 | **Untrained:** n=3 mice<br><br>**Learning:** n=5 mice | Restricted Maximum Likelihood model (REML) to predict number of dynamic sheaths depending on type of axon (untrained vs. baseline-labeled vs. learning-activated) with random variable of Mouse.<br><br>$F(2,6.68)=25.38$, $p=0.0008$<br><br>$R^2=0.94$ | All post-hoc tests conducted using Tukey's HSD:<br><br>**Untrained vs. Learning-activated:** p=0.0054<br><br>**Baseline-labeled vs. Learning-activated:** p=0.0009<br><br>All other relevant comparisons p>0.05 |
| **Fig. 6e** | Cumulative length change (in microns) of individual sheaths across time (baseline (d -21), pre-learning (d 1), directly post-learning (d 8), four weeks post-learning (d 31) on learning-activated axons compared to axons in age-matched untrained controls and baseline-labeled axons in the same mice. (mean±SEM) | **Baseline:**<br>Untrained: 0.00±0.00<br>Baseline-labeled: 0.00±0.00<br>Learning-activated: 0.00±0.00<br><br>**Pre-learning:**<br>Untrained: 1.56 ± 3.4<br>Baseline-labeled: 3.36 ± 3.08<br>Learning-activated: 1.98 ± 2.19<br><br>**Directly post-learning:**<br>Untrained: 4.33 ± 2.36<br>Baseline-labeled: -1.39 ± 0.43<br>Learning-activated: -8.14 ± 2.57<br><br>**Four weeks post-learning:**<br>Untrained: 7.69 ± 2.4<br>Baseline-labeled: 0.34 ± 0.88<br>Learning-activated: -8.32 ± 1.34 | **Untrained:** n=3 mice<br><br>**Learning:** n=5 mice | Restricted Maximum Likelihood model (REML) to predict sheath length with full factorial model between training (untrained vs. baseline-labeled vs. learning-activated) and day (baseline vs. pre-learning vs. directly post-learning vs. four weeks post-learning) and with random variable of Mouse.<br><br>Interaction effect between Training and Day to predict sheath length: $F(6,28.07)=7.14$, $p<0.0001$<br><br>$R^2=0.79$ | All post-hoc tests conducted using Tukey's HSD:<br><br>**Untrained vs. Learning-activated:**<br>d8: p = 0.0012<br>d31: p < 0.0001<br><br>**Baseline-labeled vs. learning activated:**<br>d31: p = 0.0026<br><br>**Learning-activated:**<br>**d-21 vs. d8:** p=0.0042<br>**d1 vs. d8:** p = 0.004<br>**d-21 vs. d31:** p=0.0042<br>**d1 vs. d31:** p=0.0002<br><br>All other relevant comparisons p>0.05 |
| **Fig. 6f** | Proportion of axons (%) with widening nodes, in learning-activated axons and age-matched untrained controls and baseline-labeled axons in the same mice. (mean±SEM) | **Untrained axons:** 0.00±0.00<br><br>**Baseline-labeled axons:** 28.0±11.6<br><br>**Learning-activated axons:** 76.70±14.5 | Untrained: n=4 mice<br><br>Learning: n=5 mice | Restricted Maximum Likelihood model (REML) to predict proportion of axons with widening nodes (%) depending on type of axon (untrained vs. baseline-labeled vs. learning-activated) with random variable of Mouse.<br><br>$F(2,7.97)=12.21$, $p=0.0037$<br><br>$R^2=0.83$ | All post-hoc tests conducted using Tukey's HSD:<br><br>**Untrained vs. Learning-activated:** p=0.0044<br><br>**Baseline-labeled vs. Learning-activated:** p=0.015<br><br>All other relevant comparisons p>0.05 |

| | | | | | |
|---|---|---|---|---|---|
| **Fig. 6g** | Cumulative length change (in microns) of nodes across time (baseline (d -21), pre-learning (d 1), directly post-learning (d 8), four weeks post-learning (d 31) on learning-activated axons compared to axons in age-matched untrained controls and baseline-labeled axons in the same mice. (mean±SEM) | **Baseline:**<br>Untrained: 0.00±0.00<br>Baseline-labeled: 0.00±0.00<br>Learning-activated: 0.00±0.00<br><br>**Pre-learning:**<br>Untrained: -0.92 ± 2.07<br>Baseline-labeled: 0.46 ± 0.77<br>Learning-activated: -0.61 ± 0.36<br><br>**Directly post-learning:**<br>Untrained: 0.21 ± 1.55<br>Baseline-labeled: -2.27 ± 1.41<br>Learning-activated: 7.72 ± 2.54<br><br>**Four weeks post-learning:**<br>Untrained: -2.16 ± 0.91<br>Baseline-labeled: -1.09 ± 1.74<br>Learning-activated: 8.93 ± 4.62 | **Untrained:**<br>n=4 mice<br><br>**Learning:** n=4 mice | Restricted Maximum Likelihood model (REML) to predict node length with full factorial model between training (untrained vs. baseline-labeled vs. learning-activated) and day (baseline vs. pre-learning vs. directly post-learning vs. four weeks post-learning) and with random variable of Mouse.<br><br>Interaction effect between Training and Day to predict node length: $F(6,24.32)=4.76$, $p=0.0025$<br><br>$R^2=0.78$ | All post-hoc tests conducted using Tukey's HSD:<br><br>**Untrained vs. Learning-activated:**<br>d31: p = 0.020<br><br>**Baseline-labeled vs. learning activated:**<br>d31: p = 0.047<br><br>**Learning-activated:**<br>**d1 vs. d31:** p = 0.0057<br>**d1 vs. d8:** p=0.022<br><br>All other relevant comparisons p>0.05 |
| **Fig. 6h** | Proportion of axons (%) with heightened sheath addition, in learning-activated axons and age-matched untrained controls and baseline-labeled axons in the same mice. (mean±SEM) | **Untrained axons:**<br>6.22±4.06<br><br>**Baseline-labeled axons:**<br>17.0±10.5<br><br>**Learning-activated axons:** 72.9±10.4 | Untrained: n=5 mice<br><br>Learning: n=4 mice | Restricted Maximum Likelihood model (REML) to predict proportion of axons with heightened sheath addition (%) depending on type of axon (untrained vs. baseline-labeled vs. learning-activated) with random variable of Mouse.<br><br>$F(2,6.96)=20.43$, $p=0.0012$<br><br>$R^2=0.88$ | All post-hoc tests conducted using Tukey's HSD:<br><br>**Untrained vs. Learning-activated:**<br>p=0.0016<br><br>**Baseline-labeled vs. Learning-activated:**<br>p=0.0027<br><br>All other relevant comparisons p>0.05 |
| **SUPP. FIG7** | **Measure** | **Values** | **N** | **Statistical test** | **Significance** |
| **Supp. Fig6c** | Rate of cFos+ cell gain at baseline, 2 weeks post-learning, and 4 weeks post-learning in untrained mice injected with tamoxifen, learning mice injected with tamoxifen, and learning mice injected with oil | **Baseline:**<br>**Untrained + tmx**<br>0.93±0.42<br>**Learning + tmx**<br>1.03±0.21<br>**Learning + oil**<br>1.11±0.57<br><br>**2w post-learn:**<br>**Untrained + tmx**<br>0.91±0.49<br>**Learning + tmx**<br>2.92±0.45<br>**Learning + oil**<br>0.18±0.18<br><br>**4w post-learn:**<br>**Untrained + tmx**<br>0.85±0.55<br>**Learning + tmx**<br>0.62±0.34 | **Untrained + tmx**<br>n=4 mice<br><br>**Learning + tmx**<br>n = 5 mice<br><br>**Learning + oil**<br>n = 3 mice | Restricted Maximum Likelihood model (REML) to predict rate of cell gain between training (untrained + tmx vs learning + tmx vs learning + oil) and phase (baseline vs. two weeks post-learning vs. four weeks post-learning) and with random variable of Mouse.<br><br>Interaction effect between Training and Day to predict cell gain: $F(2,16.45)=74.411$, $p=0.019$<br><br>$R^2=0.70$ | All post-hoc tests conducted using Tukey's HSD:<br><br>**Learning + tmx, baseline** vs. **Learning + tmx, 2w post-learn**: p=0.0023<br><br>**Learning + tmx, 2w post-learn** vs. **Untrained + tmx, 2w post-learn**: p=0.042<br><br>**Learning + tmx, 2w post-learn** vs. **Learning + oil, 2w post-learn**: p=0.018 |

| | | | | | |
|---|---|---|---|---|---|
| | | **Learning + oil**<br>0.27±0.19 | | | |
| **Supp. Fig6d** | Proportion of cfos+ cells gained post-learning (%) in layer 1 (L1) vs. layer 2/3 (L2/3) of cortex (mean±SEM) | **L1:**<br>14.67±8.16%<br><br>**L2/3:**<br>85.33±8.16% | n=5 learning mice | Paired student's t-test<br><br>t(4)=8.82 | p=0.010<br>Cohen's d=3.24 |
| **FIG. 7** | **Measure** | **Values** | **N** | **Statistical test** | **Significance** |
| **Fig. 7a** | Correlation between days above 20% success and axons with node lengthening (%) | | **N = 14 mice** | Linear regression between success (days and node lengthening (%)<br><br>**Baseline**<br>F(1,12) = 0.05,<br>$R^2$ = 0.004<br><br>**Learning**<br>F(1,12) = 23.57,<br>$R^2$ = 0.004<br><br>**Post-learning**<br>F(1,12) = 2.83,<br>$R^2$ = 0.191 | **Baseline**<br>p = 0.82<br><br>**Learning**<br>p = 0.7<br><br>**Post-learning**<br>p = 0.1183 |
| **Fig. 7b** | Correlation between days above 20% success and axons with sheath addition (%) | | **N = 14 mice** | Linear regression between success (days) and sheath addition (%)<br><br>**Baseline**<br>F(1,11) = 0.34,<br>$R^2$ = 0.030<br><br>**Learning**<br>F(1,11) =0.09,<br>$R^2$ = 0.008<br><br>**Post-learning**<br>F(1,12) = 8.17,<br>$R^2$ = 0.405 | **Baseline**<br>p = 0.5<br><br>**Learning**<br>p = 0.7<br><br>**Post-learning**<br>p = 0.0144 |
| **Fig. 7c** | Timing (in days) of pre-existing sheath retraction versus new sheath addition relative to the onset of learning for high performers (mean±SEM) | **Sheath retraction:**<br>6.42±1.2 days<br><br>**Sheath addition:**<br>12.00±1.09 days | n=8 learning mice | Paired student's t-test<br><br>**Learning**<br>t(5) = 3.99 | **Learning**<br>P=0.011<br>Cohen's d = 4.87 |
| **Fig. 7d** | Timing (in days) of pre-existing sheath retraction versus new sheath addition relative to the onset of learning for low performers (mean±SEM) | **Sheath retraction:**<br>9.68±1.97 days<br><br>**Sheath addition:**<br>9.2±0.881 days | n=8 learning mice | Paired student's t-test<br><br>**Learning**<br>t(4) = -0.049 | **Learning**<br>p = 0.96 |
